# bayesReact: Expression-coupled regulatory motif analysis detects microRNA activity in cancer and at the single cell level

**DOI:** 10.1101/2024.09.10.612047

**Authors:** Asta M. Rasmussen, Alexandre Bouchard-Côté, Jakob S. Pedersen

**Author notes:** The authors wish it to be known that, in their opinion, the last two authors should be regarded as Joint Last Authors.

## Abstract

**Motivation:** Regulatory constraints are crucial in maintaining tissue and cell integrity, and play important roles during developmental processes and environmental responses. Yet many regulatory mechanisms remain unobserved at the single-cell level and statistical inference may, in some cases, help elucidate their condition-specific activity and perturbation during disease progression.

**Results:** We introduce bayesReact (BAYESian modeling of Regular Expression ACTivity), a generative model of motif occurrence across experimentally ranked sequences to infer motif-based regulatory activities. The method is evaluated for microRNAs (miRNAs), which perform post-transcriptional regulation through target mRNA destabilization and translational repression. Inferred miRNA activities positively correlate with the observed miRNA expressions in primary tumors from The Cancer Genome Atlas (TCGA) and mouse stem cells. The top miRNA activity profiles are as informative for TCGA cancer-type cluster identification as the top miRNA or mRNA expression profiles. The activity captures tissue-specific mi RNA patterns observed in the matched expression, e.g., the expression of miR-122-5p in the liver and miR-124-3p in low-grade gliomas (LGG). We observe a negative association between the activity of the two miRNAs and their target gene expressions, including between the miR-124-3p activity and the anti-neuronal REST expression in LGG. bayesReact outperforms the existing method, miReact, on sparse count data, and shows a higher correlation with the miRNA expression in single-cell data. The method recovers temporal activities of prominent miRNAs during murine stem cell differentiation, including miR-298-5p, miR-92-2-5p, and the large Sfmbt2 cluster (miR-297-669). The bayesReact model is probabilistic and quantifies the uncertainty of all provided estimates. It is unsupervised and permits screens of bulk or single-cell data to identify condition-specific regulatory motif candidates. It further improves miRNA activity inference in single-cell data.

**Availability and implementation:** bayesReact is implemented as an R-package, uses a Hamiltonian Monte-Carlo sampler for posterior approximation, and is available at https://github.com/astamr/bayesReact.

## 1. Introduction

Regulatory mechanisms are vital to maintain cellular homeostasis and facilitate proper cell proliferation and differentiation while preserving tissue integrity and prohibiting carcinogenesis (Lee and Young, 2013; Slack and Chinnaiyan, 2019). However, much is still unknown regarding the spatio-temporal complexity of regulatory cell constraints, with new regulators still being discovered and characterized (Shi et al., 2022; Statello et al., 2021; Rasmussen et al., 2023). Regulatory mechanisms are frequently facilitated through motif recognition (Van Roey and Davey, 2015; Navarro et al., 2021), where a motif is a distinct biological pattern, e.g., a nucleotide (nt) or peptide sequence. Regulatory motif representations range from short strings to complex regular expressions (REs) and position weight matrices (PWMs), with examples including binding sites for transcription factors (TFs), RNA-binding proteins (RBPs), and microRNAs (miRNAs) (Van Roey and Davey, 2015; Lambert et al., 2018; Corley et al., 2020; Gebert and MacRae, 2019; Ebert and Sharp, 2012).

miRNA represents an intensely studied class of small non-coding RNA (ncRNA) with a length of 20-24 nts, which post-transcriptionally regulate target mRNAs by binding to their 3’ untranslated regions (UTRs) (Gebert and MacRae, 2019; Kilikevicius et al., 2022). Mature miRNAs originate from the 5p or 3p arm of a stem-loop precursor (Fig. 1A). In cases where both arms produce viable miRNAs, these usually differ in their seed sites and targets (Kozomara et al., 2019). The miRNA-mRNA interaction primarily occurs between the miRNA seed site, usually at nucleotide position 2-8, and a reverse complementary target site on the mRNA (Fig. 1B) (Gebert and MacRae, 2019; Agarwal et al., 2015; McGeary et al., 2019). Mature miRNA associates with the RNA-induced silencing complex (RISC) and primarily represses target mRNA translation through transcript destabilization and degradation (Guo et al., 2010; Gebert and MacRae, 2019). miRNAs modulate the abundance of their target transcripts and affect cell differentiation, proliferation, and apoptotic processes (Ebert and Sharp, 2012). Consequently, miRNAs are shown to be perturbed in cancer (Slack and Chinnaiyan, 2019; Rupaimoole and Slack, 2017; Kim and Croce, 2023), including miR-122-5p and miR-124-3p, which are found to be down-regulated in hepatocellular carcinoma (HCC) (Chun, 2022; Coulouarn et al., 2009; Ha et al., 2019) and glioblastomas (Jia et al., 2019; Silber et al., 2008), respectively. Both miRNAs are highly expressed in fully differentiated cell stages, and their down-regulation may thus promote stem-like features and subsequent cancer progression (Coulouarn et al., 2009; Silber et al., 2008). miRNAs can possess both oncogenic and tumor-suppressive capabilities (Rupaimoole and Slack, 2017), with similar trends observed for other classes of regulators (Slack and Chinnaiyan, 2019; Anastasiadou et al., 2018; Gebauer et al., 2021).

**Figure 1.**
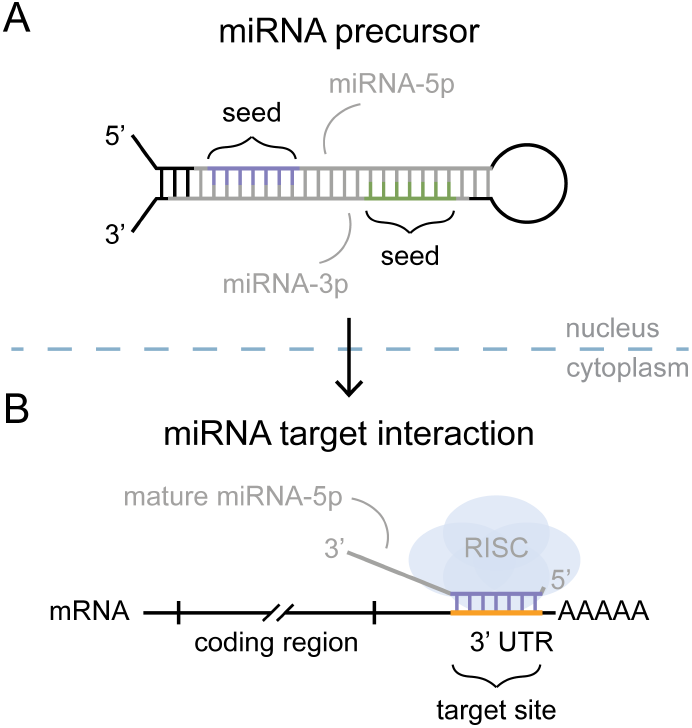
microRNA motif model and function. (A) microRNA (miRNA) precursor with core sequence elements highlighted. (B) Mature miRNA function through RNA-induced silencing complex (RISC) association and miRNA seed- and mRNA target-site interaction. UTR = untranslated region.

Despite considerable progress in understanding regulatory mechanisms, elucidating their cell-level and condition-specific activities remains restricted. For instance, commonly utilized high-throughput single-cell RNA-Sequencing (scRNA-Seq) platforms use poly(dT) primers for transcript capture and amplification, subsequently excluding most circular RNAs and small non-polyadenylated transcripts (Wang et al., 2023b; Hwang et al., 2018). Meanwhile, emerging whole-transcriptome and small ncRNA-centric scRNA-Seq methods are currently limited in their throughput, sensitivity, and ability to successfully capture both long and short transcripts (Isakova et al., 2021; Salmen et al., 2022; Li et al., 2023).

Several computational methods have been developed to indirectly predict the presence of miRNAs. Some methods leverage available paired miRNA-mRNA bulk expression data, including BIRTA, ActMiR, and miRSCAPE (Zacher et al., 2012; Fröhlich, 2015; Lee et al., 2016; Olgun et al., 2022). BIRTA jointly models TFs and miRNAs using a Bayesian regression framework, and the updated biRte allows for joint regulatory network inference based on pre-defined regulator-mRNA target interactions (Zacher et al., 2012; Fröhlich, 2015). miRSCAPE trains miRNA-dependent models on bulk data using extreme gradient boosting (Olgun et al., 2022; Chen and Guestrin, 2016). The model is subsequently used to predict miRNA expression from single-cell data in a similar condition, e.g., tissue or disease, assuming that heterogeneous bulk data translate to the single-cell level sufficiently well.

While expression measures transcript abundance, the activity provides a relative measure of the degree to which a regulator acts on its targets. Unsupervised methods, which generalize beyond miRNAs, leverage the known relationships between regulators and their targets to estimate activities. Motif-based and gene-set enrichment analysis-inspired methods offer a continuous measure of activity based on motif occurrences in experimentally ranked gene lists. For miRNAs, a shift in target-site-containing genes towards the lowly abundant end of the list indicates active miRNA presence. This approach was first implemented in Sylamer, which uses a hypergeometric statistic to evaluate over- and underrepresentation of simple nucleotide strings across ranked gene lists (Van Dongen et al., 2008). Meanwhile, cWords defines a Brownian bridge over the ranked gene list and evaluates the significance of its maximal value(Rasmussen et al., 2013; Jacobsen et al., 2010). Inspired by and extending these methods, we developed miReact (Nielsen et al., 2018; Nielsen and Pedersen, 2021), which consists of two steps: First, biases from the gene-specific nucleotide composition and sequence length are adjusted for. Second, the given motif’s correlation with gene ranks is evaluated using a modified Wilcoxon-Rank Sum test, which performed preferably compared to previous methods(Nielsen and Pedersen, 2021). Uniquely, miReact also allows for evaluating complex regular expressions, and the method was shown to capture expected miRNA activities at the single-cell level. All the methods share that their activity scores are based on p-values, comparing the observed data with a null expectation. Current methods do not explicitly model the underlying generative process driving expression-ranked motif distribution, preventing them from modeling uncertainty. Consequently, the methods do not easily extend to more complex settings, e.g., accounting for additional features such as target efficiency and integrating multiple data layers relevant to multi-omics data.

Here, we propose a generative process for motif occurrence across ranked gene lists, which we use to model motif activities and uncertainties by implementing a scalable probabilistic model in a user-friendly R package named bayesReact (BAYESian modeling of Regular Expression ACTivity). The method is demonstrated by estimating the miRNA activities from both bulk and single-cell expression data, and is found to perform better on sparse data than miReact. bayesReact permits general regulatory motif activity inference and evaluation of any regular expression; null model comparison using Bayes factors (BFs); computation of credible intervals (CIs); data simulation; and further model extensions, e.g., accounting for sequence rank uncertainty, target efficiency or pseudo-time. The model is implemented in STAN, and MCMC sampling is used for posterior approximation.

## 2. Materials and methods

### 2.1. Data collection and pre-processing

#### 2.1.1. Data Collection

Bulk expression data with matched mRNA and miRNA samples was obtained from the Cancer Genome Atlas (TCGA; n = 9,640) (Weinstein et al., 2013). The mRNA data was retrieved from the Recount3 project (Wilks et al., 2021), and the miRNA isoform data was extracted from the GDC data portal (Grossman et al., 2016). The mature miRNA read counts were evaluated separately for their 5p or 3p origins, and precursors were omitted. The miRNA expression was normalized for library size using transcripts per million (TPM) values. The final data constitutes 18,559 genes and 2,450 miRNAs expressed in primary tumor samples divided into 32 cancer types (Supplementary Table S1). To mimic scRNA-Seq read-sparsity, we simulated ten read count matrices with different levels of down-sampling. The matrices were generated by sampling mRNAs with probabilities proportional to their observed reads per kilobase of transcript per million mapped reads (RPKM) values. The RPKM values adjust for transcript length and library size. The mRNA expression was then defined as its sampling count. We obtained a whole-transcriptome scRNA-Seq dataset from Isakova et al. (Isakova et al., 2021), generated using the Smart-seq-total protocol. The dataset consists of 913 cells differentiated from primed mouse embryonic stem cells (mESCs). The cells were extracted at four different time points, with 18,900 genes expressed. We processed and annotated the data using the workflow provided by Isakova et al. (Isakova et al., 2021). miRNA entries were extracted from the whole transcriptome expression matrix, which contains joint 5p and 3p arm expression based on the genomic origin of the stem-loop precursor. The miRNAs were annotated using miRBase (Kozomara et al., 2019) and assigned either the 5p or 3p target site based on the overall correlation with their inferred activities.

We extracted all human 3’ UTR sequences as provided by GENCODE v. 32 and mouse sequences from GENCODE M23 (Frankish et al., 2021). The sequences were filtered to include instances with a length between 20–10,000 nts, and only the longest 3’ UTR isoform for each protein-coding gene was retained. We represent RNA sequences, transcripts, and motifs by the complementary DNA (cDNA), corresponding to replacing uracil (U) with thymine (T).

miRBase v. 22 was used for all miRNA annotations and for extracting seed sites (Kozomara et al., 2019). The 2,450 miRNAs expressed in the TCGA data shared 1,941 unique seed sites.

#### 2.1.2. Data pre-processing

The input data for bayesReact consists of three data types: first, a numerical experimental readout such as read-counts from an RNA-Seq experiment; second, a list of sequences; third, a set of motifs that interact with a subset of the sequences and are expected to affect the readout when present. During this study, miRNA-guided depletion of target transcripts is used to evaluate the performance of bayesReact. Here, the readouts represent gene expression, the sequences are 3’ UTRs, and the motifs are miRNA target sites (Supplementary Fig. S1).

##### Expression data

The initial gene expression matrix can be provided with either normalized or raw counts, which are subsequently re-scaled and log_2_-transformed using the function bayesReact::norm_scale_seq() (Supplementary Fig. S2). For raw data, the function normalizes the expression data with a pseudo-library size of summed read counts in each condition (cell, sample, or other experimental condition), which scales the expression to sum to one. Each entry is subsequently multiplied by the median expression of the condition and log_2_-transformed. This normalization procedure, called log_2_ pseudo-TPM, was used on all mRNA count data.

##### Sequence ranking

The sequences were matched to the gene identifiers of the expression matrix, and expression levels were used as proxies for sequence abundances. Fold-change (FC) scores were calculated as the log-transformed expression of gene *i* under condition *c* subtracted the gene expression from a control setting. As a default, we represent the control setting by a pseudo-normal condition due to the frequent lack of control samples, e.g., lack of healthy tissue samples matching tumor biopsies. The pseudo-normal condition is defined by the median gene expressions across all conditions in a dataset. Subsequently, tissue-specific expression patterns are assumed to drive deviations from the pseudo-normal for the TCGA data. The FC-scores were used to perform condition-specific sequence ranking, and sequences are ranked in decreasing order (Fig. 2A).

**Figure 2.**
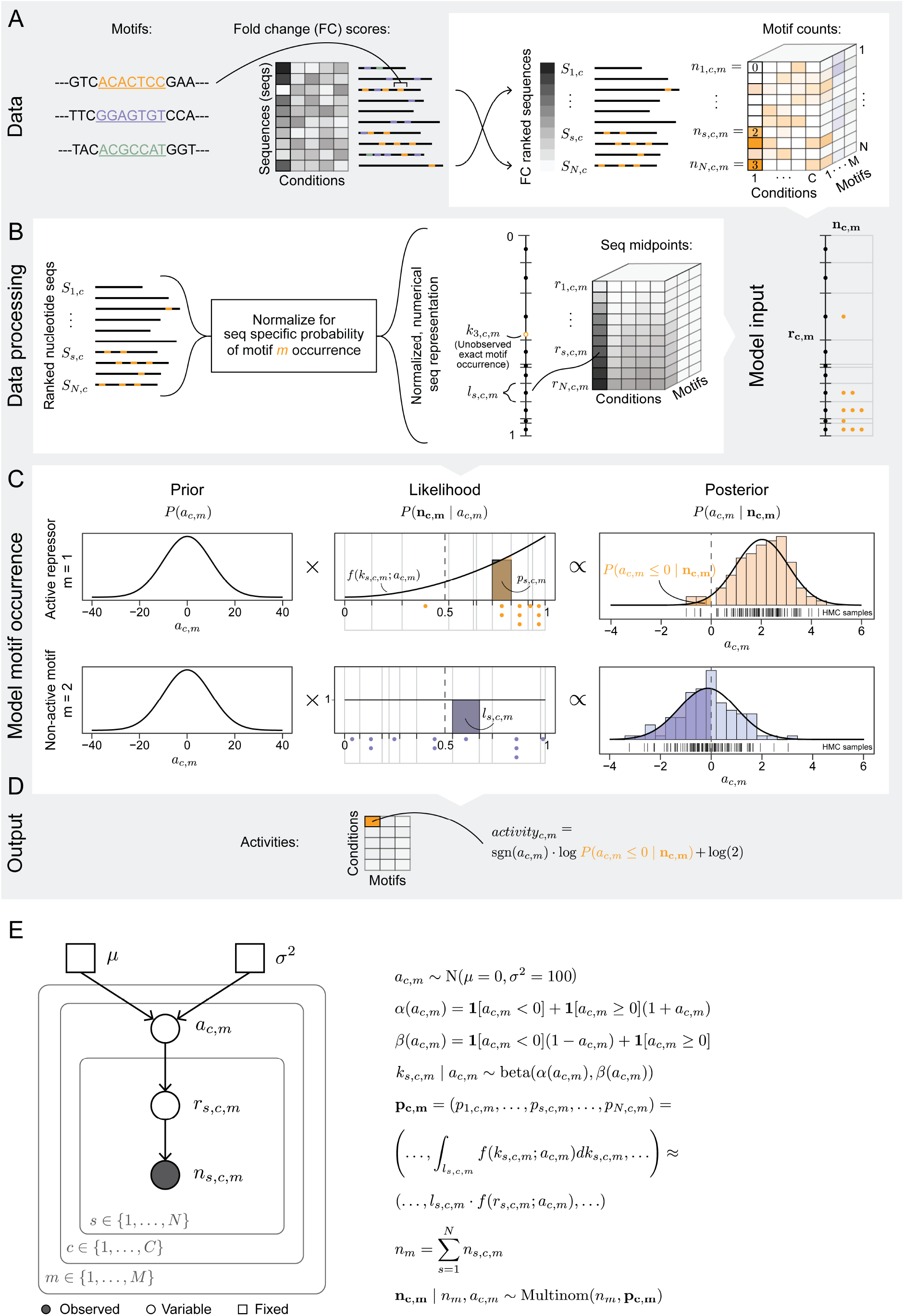
Illustration of data processing and the model framework. All necessary input data (grey background), data processing, and motif modeling (white background) are depicted. (A) Overview of bayesReact input data, with an example including 7-mer motifs, a small simulated data set of fold-change (FC) scores across multiple conditions, and sequences with annotated motifs. To the right is depicted FC-based sequence ranking and motif distribution for a single annotated motif. *S*_*s,c*_ annotates the sequence with rank *s* in condition *c*, while *n*_*s,c,m*_ the corresponding number of motif occurrences of motif *m*. (B) Sequence normalization is performed to adjust for sequence length and nucleotide composition bias, and are scaled such that the combined sequence length sums to one. *S*_*s,c*_ is then represented by a numerical motif-dependent length *l*_*s,c,m*_ and its rank *s*, which specifies the sequence location along the combined sequence interval [0, 1]. Exact motif occurrence on the combined normalized and rescaled sequence interval [0, 1] is a latent variable *k*_*s,c,m*_, and each of the *n*_*s,c,m*_ motif occurrences are represented by the sequence midpoint *r*_*s,c,m*_ (left). (C) Setup for inferring motif activities based on modeling motif occurrences across the ranked sequences. The motif distributions are parameterized by an underlying activity parameter *a*_*c,m*_. The examples shown include an active repressor (top) and an inactive or non-functional motif (bottom). The occurrences of the first motif are sequence rank-correlated, while uncorrelated for the second motif. (D) The inferred motif activities for each motif *m* in each condition *c* are output. The activity (score) is the signed log posterior probability of having parameter values with the opposite sign of the posterior mean. (E) Graphical plate representation of the bayesReact model and its dependency structure. *a*_*c,m*_ is the underlying activity parameter for motif *m* in condition *c*; *k*_*s,c,m*_ is the latent motif occurrence on the normalized ranked sequence interval [0, 1]; *p*_*s,c,m*_ is the probability of motif *m* occurrence on *S*_*s,c*_ under *f* (*k*_*s,c,m*_; *a*_*c,m*_); *n*_*m*_ the total number of motif occurrences; **n**_**c**,**m**_ the motif count distribution across all sequences. Edges indicate dependencies, squares are fixed parameters, white circles are free parameters, and the grey circle is the observed count data.

##### Motif probabilities

A motif refers to any regular expression (RE) on the alphabet {*A, T, G, C*}. In this study, we primarily focus on the set of all 7-mers (*M* = 16,384), of which a subset are miRNA targets.

The sequence-specific probability (SSP) of observing motif *m* at least once in a random sequence with length and nucleotide composition given by sequence *i* (from gene *i*) was computed using the Regmex package (Nielsen and Pedersen, 2021). Briefly, stochastic motif generation is modeled by an absorbing Markov chain, where nucleotides are drawn with probability equal to their frequency in the sequence *i* upon transition across the state-space defined by the motif RE.

### 2.2. Modeling motif activity

We developed a Bayesian method, bayesReact, to evaluate the association of a motif with the gene expression pattern and, hence, the ranking of sequences for a given condition. bayesReact models the activity of functional motifs based on motif occurrence across FC-ranked sequences. Using a generative, probabilistic approach facilitates model extensions and allows the uncertainty of activity estimates to be quantified.

#### 2.2.1. Data representation and null expectation

Let {*S*_1,*c*_, *S*_2,*c*_, …, *S*_*s,c*_, …, *S*_*N,c*_} be a set of FC-score ranked sequences, where index *s* ϵ {1, …, *N* } defines the rank, *c* ϵ {1, …, *C*} the condition, and let *m* ϵ {1, …, *M* } index the set of *M* independent motifs (Fig. 2). The ranked sequences are arranged consecutively to represent non-overlapping intervals in a given condition *c*, and the number of motif occurrences in each sequence, for a motif *m*, are considered independently Poisson-distributed random variables *n*_*s,c,m*_ ∼ Pois(*λ*_*s,c,m*_). From the sequence-specific probability, it is possible to define the Poisson rate parameter *λ*_*s,c,m*_ as follows: 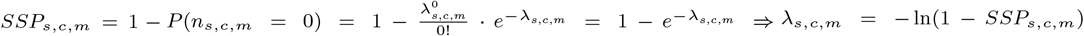 We can thus represent each *S*_*s,c*_ as a sub-interval with width corresponding to the expected number of motif occurrences given by *λ*_*s,c,m*_. This normalizes for the probability of observing *m* given the sequence length and nucleotide composition of *S*_*s,c*_. Consequently, the total number of motif counts can also be described through a Poisson process, 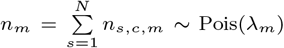 Pois(*λ*_*m*_), where 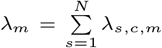 is the length of the combined sequence interval [0, *λ*_*m*_], which represents the total expected number of motif occurrences. Conditioning on *n*_*m*_, we can describe the motif count observations jointly as **n**_**c**,**m**_ = (*n*_1,*c,m*_, …, *n*_*s,c,m*_, …, *n*_*N,c,m*_) | *n*_*m*_ ∼ Multinom(*n*_*m*_, **l**_**c**,**m**_ = (*l*_1,*c,m*_, …, *l*_*s,c,m*_, …, *l*_*N,c,m*_)), where 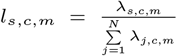 and **l**_**c**,**m**_ is a vector of probabilities. Here *l*_*s,c,m*_ is the normalized and re-scaled representation of *S*_*s,c*_ in relation to motif *m*, such that 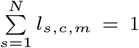 (Fig. 2B). Motif occurrence, *k*_*s,c,m*_, can be described as a stochastic process along the re-scaled sequence interval. The exact motif position on [0, 1] is unobserved and considered an auxiliary latent variable. Given the data processing and representation, the null model of no motif activity is a uniform distribution of motif events across the ranked and normalized sequences. The null model is thus:

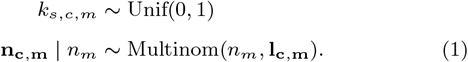

Consequently, non-functional motif events are expected to occur with equal probability across [0, 1], when we account for sequence length and nucleotide composition bias. The null model for the number of motif occurrences *n*_*s,c,m*_ in a sequence *S*_*s,c,m*_ depends only on the length of its sub-interval *l*_*s,c,m*_.

#### 2.2.2. Modeling motif occurrence and number of motif events across ranked sequences

Deviations from the null model described above can, for example, arise when a microRNA acts on a set of target transcripts. The depleted target sequences will be systematically skewed toward the end of the ranked sequence list (Fig. 2C). This signal can be captured by letting the motif occurrence be distributed according to a flexible beta distribution with support on [0, 1]. The null model then becomes a special case, 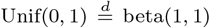, and the probability density function (PDF; *f* (*k*_*s,c,m*_; *a*_*c,m*_)) is given by:

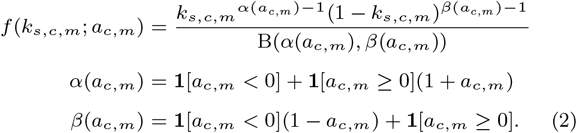

The beta shape parameters are transformations of an underlying activity parameter *a*_*c,m*_ ϵ R. The activity parameter has a clean interpretation concerning motif distribution; *a*_*c,m*_ < 0 entails motif clustering at the beginning of the cumulative sequence interval constituting sequences with high relative gene expression (FC-scores); *a*_*c,m*_ = 0 corresponds to the null model; and *a*_*c,m*_ > 0 implies motif over-representation at the end of [0, 1] for sequences with low relative abundance (Fig. 2C).

Under the beta distribution, we can describe the probability of motif occurrence on 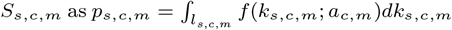 where **p**_**c**,**m**_ = (*p*_1,*c,m*_, …, *p*_*s,c,m*_, …, *p*_*N,c,m*_) is a vector of motif probabilities. The number of motif occurrences across the sequences can subsequently be described by a conditional multinomial distribution:

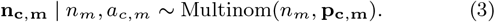

Defining **p**_**c**,**m**_ through a beta distribution, compared to a generic Dirichlet distribution, constrains the probability vector to account for sequence ranking. While the integral 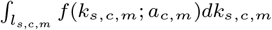 can be computed through the cumulative distribution function (CDF; *F* (.)), this is computationally intensive for large data sizes. Instead, we approximate the beta distribution with a step-function, where 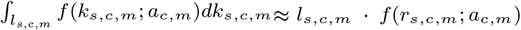 and *r*_*s,c,m*_ is the mid-point of the normalized sequence interval for *S*_*s,c,m*_ (Fig. 2B). Increasing the number of sequences *N*, leads to a finer sequence-based partitioning of [0, 1], e.g., using all human 3’ UTRs divides the cumulative sequence interval into ∼20K sub-intervals. For all *k*_*s,c,m*_ on *l*_*s,c,m*_, we have that 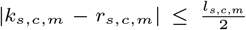 and we approximate *f* (*k*_*s,c,m*_; *a*_*c,m*_) ≈ *f* (*r*_*s,c,m*_; *a*_*c,m*_).

Subsequently, the joint probability of the observed motif counts across all sequences, conditional on the underlying activity parameter, can be approximated by the following parameterization of the multinomial probability mass function (PMF):

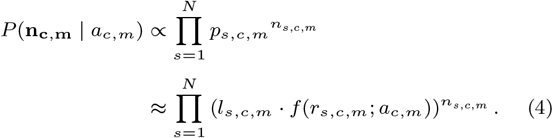

An additional benefit of approximating **p**_**c**,**m**_ is the ability to pre-compute part of the log-likelihood (gray underlining), allowing for further computational speed-up:

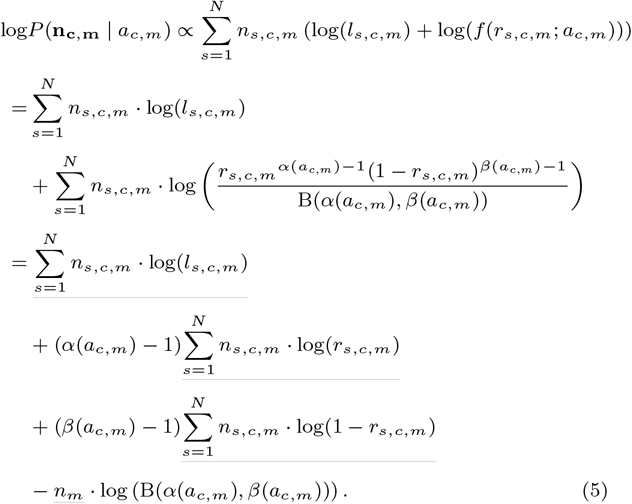

Finally, we set an uninformative prior for the activity parameter centered at zero: *a*_*c,m*_ ∼ N(*µ* = 0, *σ*^2^ = 100), where *σ* is the standard deviation.

By establishing the log-likelihood and prior distribution of *a*_*c,m*_, it is possible to explore the marginal posterior density of interest, after marginalizing the latent variable *k*_*s,c,m*_:

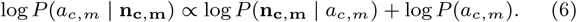

Markov chain Monte-Carlo (MCMC) sampling is used to sample from the posterior target distribution. After an approximation of the posterior is obtained, we find the activity score of motif *m* in condition *c* based on the posterior probability of the signed activity parameter being smaller or equal to zero (corresponding to the probability of no motif activity after observing the data; Fig. 2D):

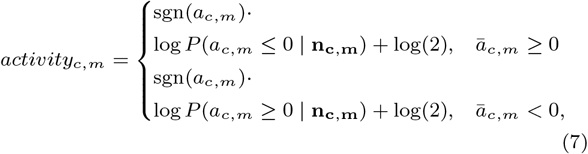

where *ā*_*c,m*_ is the posterior mean of the marginal density *P* (*a*_*c,m*_ | **n**_**c**,**m**_). The activity score constitutes two posterior probabilities and to avoid discontinuity, all values have been added a constant *log*(2).

The use of MCMC for activity inference permits modeling flexibility. However, even for a large number of MCMC samples, the tails of the posterior distributions are not efficiently characterized, and normal approximations of the MCMC samples are used instead, motivated by the Bernstein-von Mises theorem (asymptotic normality of Bayesian posteriors for large *n*_*m*_, see, e.g., Van der Vaart (1998)). The normal approximation has minimal influence on results with little or no activity. However, for motifs with substantial association with sequence ranking, the approximation avoids issues with *P* (*a*_*c,m*_ ≤ 0 | **n**_**c**,**m**_) evaluating to zero, and we maintain resolution for highly active motifs.

Establishing a generative model enables exploring the underlying processes driving motif occurrence and clustering across experimentally ranked sequences of interest (Fig. 2E). Here, motif occurrence is described under a beta distribution and controlled by an underlying activity parameter whose prior and posterior densities are both approximately normal. The activity measure captures how skewed the motif distribution is across the ranked sequences and, equivalently, the effect of a regulator on its target motifs.

#### 2.2.3. Bayes factor for null model comparison

A motif occurrence can be either non-functional (generated under a uniform distribution) or functional (generated under a beta distribution). In general, we expect occurrences of *m* on the set of sequences to be a combination of true functional motif occurrence events (a subset of which is targeted in a given condition *c*) and non-functional false positives. The null (*M*_0_; *a*_*c,m*_ = 0) and alternative (*M*_1_) models can be compared based on their marginal log-likelihoods using Bayes factor (BF):

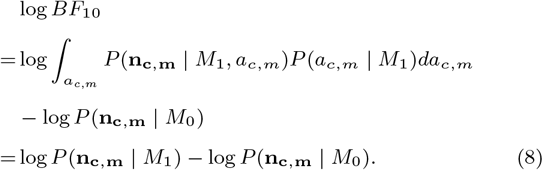

Due to the direct comparison of marginal log-likelihoods, the normalizing constants cannot be disregarded and are instead approximated using bridge sampling (Gronau et al., 2017). The BF is useful to evaluate how well *M*_1_ describes the motif observations compared to *M*_0_, particularly for conditions with activities deviating slightly from zero.

#### 2.2.4. Flexible two-parameter beta model

A more flexible model was also implemented and evaluated, where the two beta parameters *α*_*c,m*_ and *β*_*c,m*_ are freely variable instead of transformations of *a*_*c,m*_. The two-parameter beta model and corresponding activity are defined as follows:

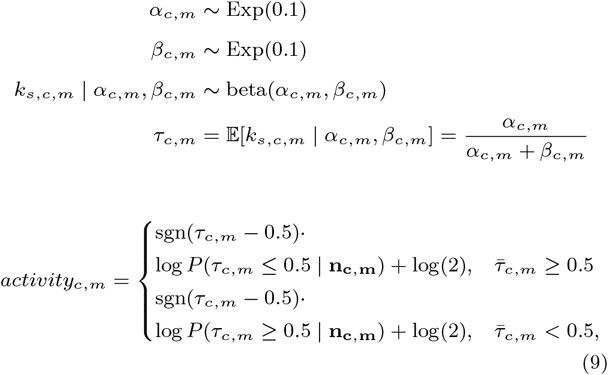

where 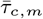 is the mean value of the marginal posterior density *P* (*τ*_*c,m*_ | **n**_**c**,**m**_).

We refer to the two models as bayesReact and bayesReact_2p_, respectively.

### 2.3. bayesReact implementation

bayesReact is an R package implemented in R, Bash, and STAN (through RSTAN (Stan Development Team et al., 2020)), allowing for user-defined evaluation of motif activities across experimentally ranked sequences. The package constitutes three primary modules: Data pre-processing, modeling motif activities and posterior approximation, and parallelization (Supplementary Figure S2). STAN’s Hamiltonian Monte Carlo (HMC) algorithm is used for MCMC sampling (Carpenter et al., 2017), and bayesReact can return the complete set of posterior samples, summary statistics such as the posterior mean of *a*_*c,m*_, credible intervals (CIs) and model diagnostics, or simply return the activity scores directly. To avoid large influences of individual sequences on the distribution of motif occurrences, a user-defined threshold can be placed on *l*_*s,c,m*_ and *n*_*s,c,m*_, with a default of 10^−6^ and 2, respectively. The user can also specify the number of MCMC chains, the number of MCMC samples, and whether to compute BFs.

## 3. Results

### 3.1. Model performance on bulk pan-cancer data

Initial model evaluation on the TCGA data showed a rapid convergence rate, with the model usually converging on the stationary distribution (supposed true posterior) within a hundred iterations. We subsequently set the default number of MCMC iterations to 3,000 and the warm-up period of 500 iterations to be discarded. Running three independent MCMC chains for all 7-mer motifs across the pan-cancer primary tumor samples shows reliable convergence 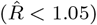 for 99.99% of the activity parameters *a*_*c,m*_ (11 · 10^3^ of 158 · 10^6^ parameters have 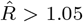 Supplementary Fig. S3A). Additionally, the effective sample sizes (ESSs) tend to be large, indicating low auto-correlation and efficient exploration of the stationary posterior distribution (99.85% of the activity parameters have an ESS of more than 1,000; Supplementary Fig. S3B). Rerunning bayesReact and increasing the number of iterations showed improved diagnostics for individual cases.

Comparable results are also observed for the more flexible bayesReact_2p_ model, with the freely variable *α*_*c,m*_ and *β*_*c,m*_ parameters (100% of 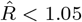 and *ESS >* 1, 000; Supplementary Fig. S3C). bayesReact and bayesReact_2p_ produce similar activity estimates and correlation to the observed miRNA expression data (see Fig. 3). bayesReact obtains a mean activity score of 0.03 and standard deviation (sd) of 3.12 across all 7-mer motifs and tumor samples. Similarly, bayesReact_2p_ provides a mean of 0.02 (sd = 3.29). The low mean activity is consistent with the majority of 7-mers expected to be non-functional and that many functional regulatory motifs are tissue and cancer-type specific. Due to the comparable results between the two models, we proceed using the model with the fewest free parameters as the core of bayesReact.

**Figure 3.**
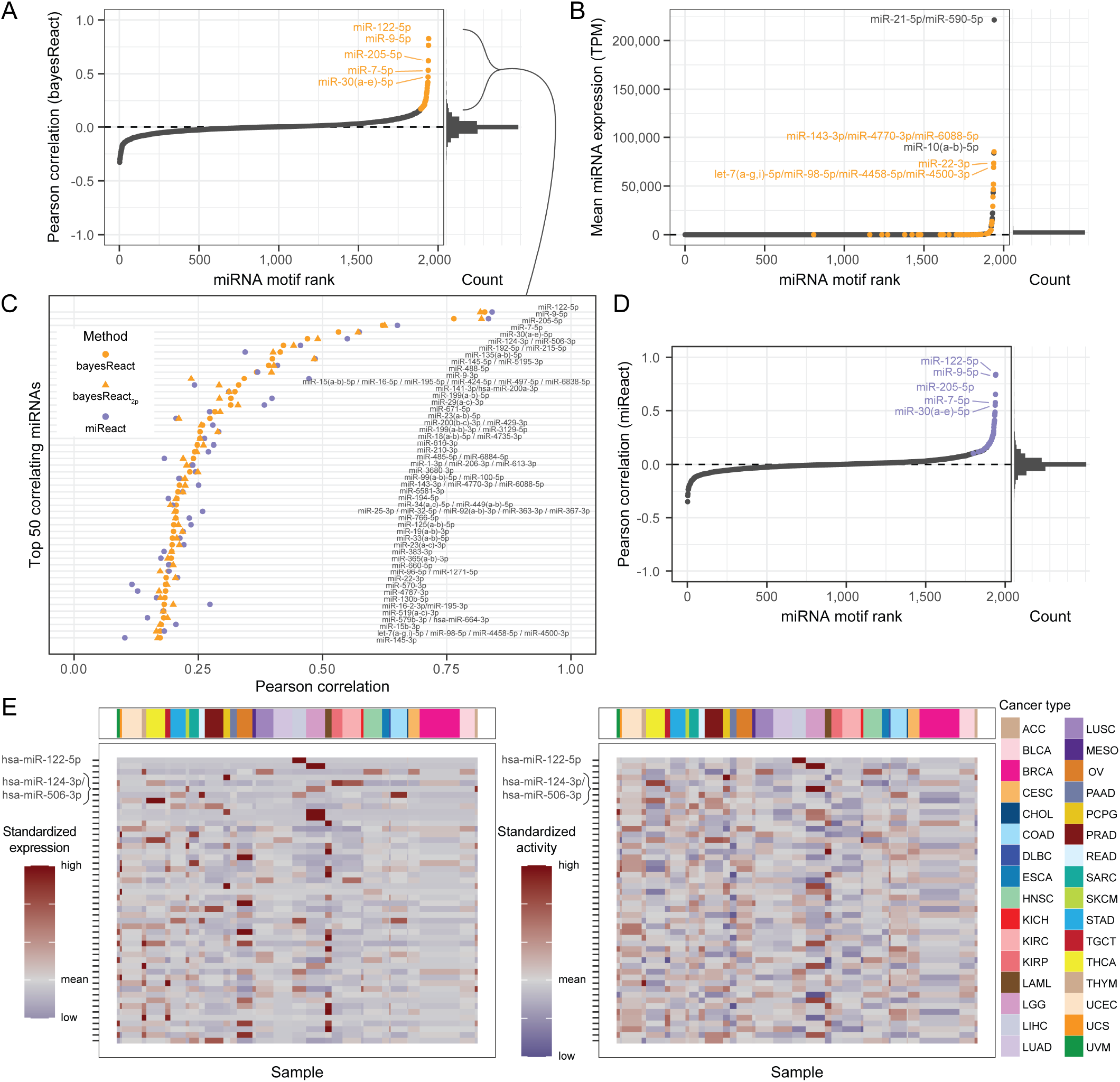
Pan-cancer microRNA activity and expression overview. (A) Pearson correlation between miRNA expression and activity across all primary tumor samples (left). Depicted are 2,450 miRNAs with expression collapsed by their shared target sites (n = 1,941). The miRNAs and their target motifs are subsequently ranked by the correlation. The top 50 miRNAs with the highest correlation coefficients are highlighted in orange, and the top 5 are annotated. On the right is depicted the corresponding histogram containing 50 bins. (B) miRNAs ranked by average expression across all TCGA samples (left). The top 50 correlating miRNAs from panel A are highlighted in orange, and the top 5 are annotated. On the right is depicted the corresponding histogram with 50 bins. TPM = transcripts per million. (C) Top 50 miRNAs based on activity and expression correlation across all samples. The Pearson correlation is shown for three different methods of activity inference: bayesReact, bayesReact_2p_, and miReact. (D) Pearson correlation between miRNA expression and activity scores obtained through miReact. miRNA target motifs are ranked by their correlation coefficient. The top 50 miRNAs from panel A are highlighted in purple, and the top 5 are annotated. (E) Heatmaps for the top 50 correlating miRNAs ordered by correlation coefficient and clustered by cancer type. The mean expression for each cancer type (left) and bayesReact activity (right) are depicted. Values are standardized for visualization purposes.

Comprehensive model evaluation was performed for individual *a*_*c,m*_ (Supplementary Fig. S3D-G), with prior and posterior predictive checks showing vastly improved agreement between observed cumulative motif distributions and simulated motif data under the posterior predictive distribution compared to the prior predictive distribution (Supplementary Fig. S3E,G).

Finally, we evaluated the variability in activity estimation, a product of stochastic MCMC sampling. This was done by conducting 1,000 bayesReact repetitions for the two motifs ’ACACTCC’ (miR-122-5p target) and ’GTGCCTT’ (miR-124-3p target) across all TCGA samples. Each repetition was summarised by the activity correlation with the observed miRNA expression data. Encouragingly, the correlation showed low variability, and the coefficients range from 0.824 − 0.828 and 0.456 − 0.464, respectively, and the median and mean values coincide in both instances (Supplementary Fig. S3H). Equivalent results are observed for bayesReact_2p_.

#### 3.1.1. microRNA activity in cancer

The miRNA expression is generally small across the pan-cancer TCGA data with an overall mean expression of 504.73 TPM (sd = 9,277.21 and median = 0 TPM), and the majority of miRNAs having a mean expression across the tumor samples close to zero (Fig. 3B). The TCGA data comprises 9,640 primary tumor samples categorized into 32 cancer types originating from 27 distinct tissues (Supplementary Table S1). Performing hierarchical clustering, with the number of clusters predefined as the number of cancer types (n = 32), we find that the mRNAs and miRNAs with the highest mean expression (n = 100) recover the cancer-type clusters to the same degree. The clustering yields adjusted rand indexes (ARIs; (Scrucca et al., 2023)) of 0.20 and 0.19, respectively (Supplementary Fig. S4A-B). Interestingly, the corresponding inferred miRNA activities recovered the cancer clusters equally well, indicating the same level of information content present to differentiate between cancer types (ARI = 0.22; Supplementary Fig. S4C).

While the activity of a miRNA does not directly correspond to its expression level, the two variables are still expected to be associated. An elevated cytoplasmic miRNA content can increase the degradation of its mRNA targets, while non-transcribed miRNAs are inactive (Gebert and MacRae, 2019). We exploit this relationship to evaluate the performance of bayesReact using the correlation between the observed miRNA expression and inferred activity. The mean Pearson correlation (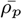; sensitive to tissue-specific outliers) is small for the set of 2,450 expressed miRNAs collapsed by their 1,941 shared target motifs (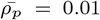; Fig. 3A). This is unsurprising given the miRNAs’ overall limited expression and activity across the primary tumors. Similar results are also obtained using the existing method miReact (Nielsen and Pedersen, 2021) to estimate miRNA activities (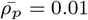; Fig. 3C-D), with the method applied to the same input data as bayesReact for comparative analysis. Both methods tend to recover the same miRNAs with large correlation coefficients, suggesting that they recover the same underlying motif distribution signal from the bulk expression data (Fig. 3A,C-D; Supplementary Table S2).

We generally observe the top correlating miRNAs to have a tissue-specific expression and activity pattern (Fig. 3E; Supplementary Table S2). Prominent examples include miR-122-5p (*ρ*_*p*_ = 0.83), primarily expressed in the liver, as well as miR-9-5p (*ρ*_*p*_ = 0.76) and miR-124-3p (*ρ*_*p*_ = 0.46) expressed in the brain (Fig. 3C,E).

miR-122-5p activity in liver hepatocellular carcinoma miR-122-5p, which has the highest association between expression and activity, is known to be highly expressed in liver tissue (Lagos-Quintana et al., 2002) and is predominantly expressed in the liver hepatocellular carcinoma samples (LIHC; n = 367; Fig. 4A). The expressed miR-122-5p is expected to deplete its target transcripts actively, and concordantly, we observe a clear association between the occurrence of the miR-122-5p target motif in the 3’ UTR of a gene and its relative expression (position along the FC-ranked 3’ UTR sequence interval; Fig. 4B). In contrast, the complementary miR-122-5p seed site distribution across the FC-ranked sequences is uniform, as it does not represent a regulatory motif in the 3’ UTRs (Fig. 4B). Using the observed target motif distributions (**n**_**c**,**m**_) and MCMC sampling, we recovered the marginal posterior distributions for the miR-122-5p activity parameters (n = 367). Summarising the distributions by the posterior mean and 99% CI, we find that the credible intervals mostly overlap zero for LIHC samples with low miRNA expression (Supplementary Fig. S5A). Subsequently, the joint posterior probability **1**[*ā*_*c,m*_ ≥ 0] log *P* (*a*_*c,m*_ ≤ 0 | **n**_**c**,**m**_) + **1**[*ā*_*c,m*_ < 0] log *P* (*a*_*c,m*_ c 0 | **n**_**c**,**m**_) + log(2) (from eq. 7) is larger in liver cancer compared to other cancer types (Supplementary Fig. S5B), resulting in a LIHC-specific miR-122-5p activity. Furthermore, TCGA samples with activity scores close to zero also tend to have log *BF*_10_ ≤ 0, indicating no support for bayesReact compared to the uniform null model (Fig. 4C). Concludingly, bayesReact efficiently recovers condition-specific miR-122-5p activities, which agree with the observed miRNA expression and known liver-specific function. The results are consistent with previous miReact findings (Nielsen and Pedersen, 2021) and showcase bayesReact’s extended capabilities in quantifying uncertainties for the underlying activity parameters and subsequent regulatory motifs of interest.

**Figure 4.**
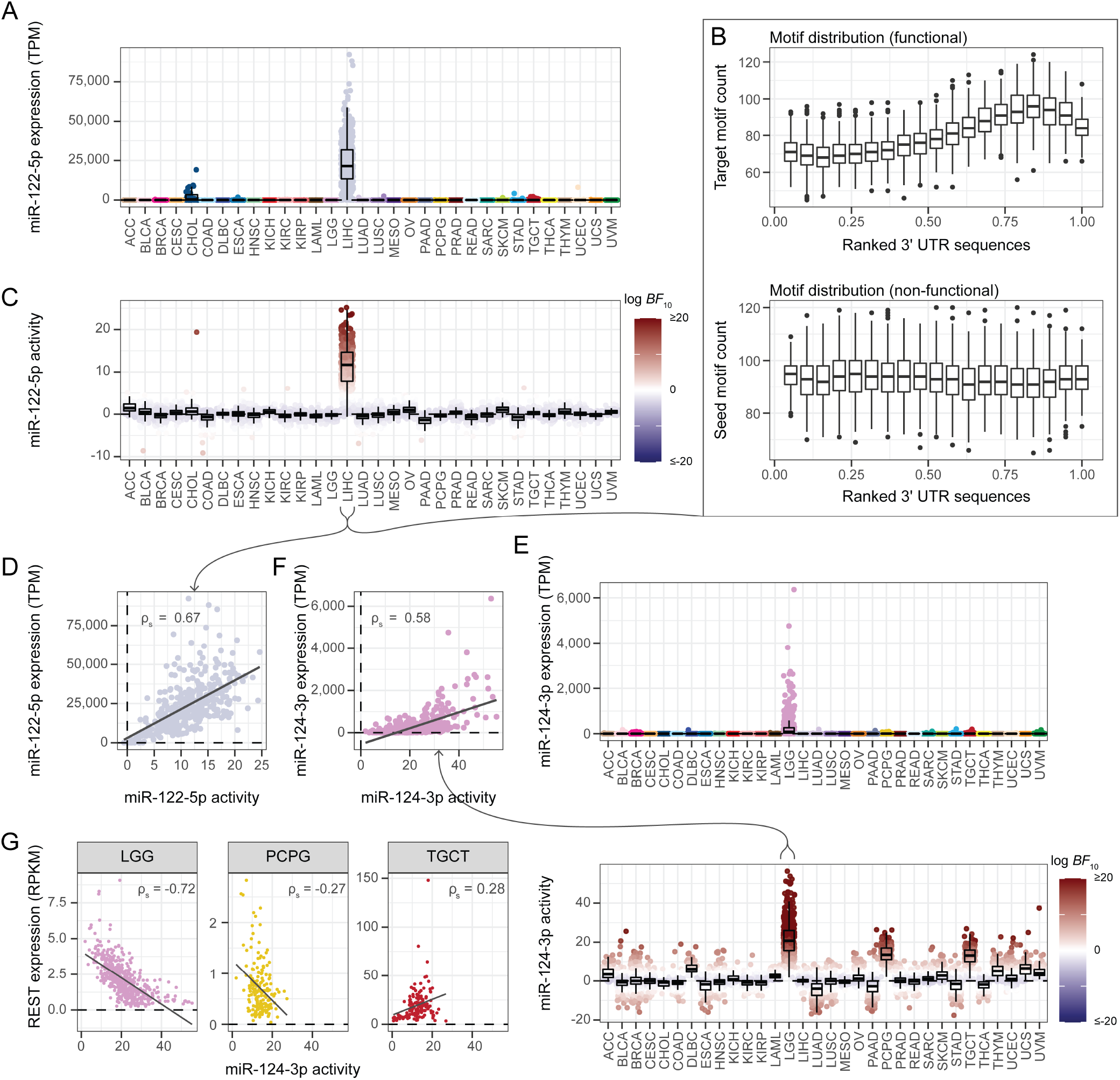
Tissue-specific miR-122-5p and miR-124-3p activities. (A) miR-122-5p expression for each primary tumor sample grouped by cancer type. TPM = transcripts per million. (B) miR-122-5p target site (top) and its complementary seed site (bottom) distribution across the normalized 3’ untranslated region (UTR) sequences. The combined 3’ UTR sequence interval is divided into 20 bins, and each boxplot depicts the motif count within a bin for each LIHC sample (n = 367). The functional miR-122-5p target motif occurs in the 3’ UTRs and is evaluated to infer the miRNA activity (see Fig. 1B). The target motif occurs 1,692 times across 1,526 3’ UTRs, while the seed motif occurs 1,978 times in 1,768 3’ UTRs. (C) miR-122-5p activities grouped by cancer type. Each sample is annotated with the log *BF*_10_ value. BF = Bayes factor. (D) Scatterplot of miR-122-5p activity against expression, with the Spearman correlation (*ρ*_*s*_) annotated and linear regression line shown. (E) miR-124-3p expression (top) and activity (bottom) across all TCGA samples. Top: Samples are annotated by cancer type. Bottom: Samples are annotated by their log *BF*_10_ values. (F) miR-124-3p activity plotted against the observed expression, with the Spearman correlation annotated and linear regression line shown. (G) Association between the miR-124-3p activity and the expression of its downstream target REST in low-grade gliomas (LGG; n = 509; left), pheochromocytoma and paragangliomas (PCPG; n = 182; middle), and testicular germ cell tumors (TGCT; n = 139; right). Expression values are provided as reads per kilobase of transcript per million mapped reads (RPKM). Spearman correlations (*ρ*_*s*_) and linear regression lines are visualized.

Considering the inferred LIHC activities, a positive Spearman correlation (*ρ*_*s*_; robust to outliers) is observed between the miR-122-5p activity and expression (*ρ*_*s*_ = 0.67; Fig. 4D). In comparison, *ρ*_*s*_ = 0.64 for the miReact activities. In addition, the activity negatively correlates with the expression levels of the known target genes Rac1 and RhoA (*ρ*_*s*_ ≤ −0.33; Supplementary Fig. S5C; (Wang et al., 2014)), and a positive association is observed between the miRNA activity and the TF promoting its host gene transcription (*ρ*_*s*_ = 0.42; Supplementary Fig. S5D; (Li et al., 2011)). The observed miR-122-5p expression showed similar associations with the expression of the target genes (*ρ*_*s*_ ≤ −0.38) and TF (*ρ*_*s*_ = 0.39).

The miR-124-3p activity negatively associates with the expression of the anti-neuronal RE1-silencing transcription factor miR-124-3p promotes a neuronal cell fate for differentiating neuroectodermal progenitors (Makeyev et al., 2007; Stark et al., 2005), and is expressed in low-grade gliomas (LGG) as well as a moderately expressed in pheochromocytomas and paragangliomas (PCPG) and testicular germ-cell tumors (TGCT; Fig. 4E, Supplementary Fig. S5E). A positive correlation is observed between the miR-124-3p expression and activity across all cancer types (Fig. 3C) and within cancer types with significant miR-124-3p expression (*ρ*_*s*_ ≥ 0.26; Fig. 4F, Supplementary Fig. S5F-G). The LGG samples showed the largest expression, activity, and subsequent association between the two (*ρ*_*s*_ = 0.58 using bayesReact and *ρ*_*s*_ = 0.57 with miReact).

miR-124-3p partake in a double-negative feedback loop by antagonizing the RE1-silencing transcription factor (REST)/Scp1 pathway (Visvanathan et al., 2007; Conaco et al., 2006; Stappert et al., 2015; Suster and Feng, 2021). miR-124-3p represses Scp1, which is observed to be almost completely depleted in the LGG samples (Supplementary Fig. S5H), and is in turn expected to destabilize the REST protein (Burkholder et al., 2018), which otherwise enables the suppression of neuron-specific gene transcription and promote non-neuronal cell states (Visvanathan et al., 2007; Burkholder et al., 2018). We observe a strong negative correlation between the inferred miR-124-3p activity and REST expression in LGG samples (*ρ*_*s*_ = −0.72; Fig. 4G), which, intriguingly, is larger than for the corresponding miRNA expression (*ρ*_*s*_ = −0.32; Supplementary Fig. S5H), and a negative correlation is only observed in tumors originating from the central nervous system (CNS; Fig. 4G).

Interestingly, miR-9-5p, known to target the REST transcript directly (Packer et al., 2008; Stappert et al., 2015; Godlewski et al., 2019), showed a smaller correlation with the REST expression (*ρ*_*s*_ = −0.33 for its activity and *ρ*_*s*_ = −0.31 based on the expression). The results encourage further investigation of additional regulatory mechanisms bypassing Scp1, e.g., the direct targeting by miR-124-3p of the REST complex transcripts, in line with previous observations of potential miR-124-3p target sites present in 3’ UTR of the CoREST and MeCP2 genes (Wu and Xie, 2006; Baudet et al., 2012). Alternative candidates are PTBP1 or BAF53a, which are also part of the regulatory REST circuit and have previously been shown to be depleted by miR-124-3p (Makeyev et al., 2007; Yoo et al., 2009; Stappert et al., 2015; Suster and Feng, 2021). The expression of both PTBP1 and BAF53a correlate negatively with the miR-124-3p activity in low-grade gliomas (*ρ*_*s*_ = −0.44 and *ρ*_*s*_ = −0.55, respectively) and to a lesser degree with the miRNA expression (*ρ*_*s*_ = −0.42 and *ρ*_*s*_ = −0.40, respectively).

#### 3.1.2. microRNA activity inference on sparse expression data

While miRNA expression can be recovered from small RNA-Seq protocols, it is not recovered when performing high-throughput scRNA-Seq. Activity inference thus presents a unique opportunity to indirectly investigate the cell-level miRNA activity. To evaluate the performance of bayesReact on data with similar read sparsity as scRNA-Seq data, we generated ten TCGA expression matrices with varying library sizes (total number of gene counts per sample; see subsection 2.1.1), which were used as input for the miRNA activity inference by bayesReact and miReact. In general, increased read count sparsity led to decreased activity scores and subsequent lower correlation between the miRNA activity and its expression (Fig. 4A). Focusing on the top 50 correlating miRNAs (Supplementary Table S2), bayesReact tends to retain a higher correlation score with increasing count sparsity than miReact (Fig. 4A-B). miReact recovers the activity slightly better for large library sizes (> 500, 000 gene counts), while bayesReact outperforms miReact at count sparsities resembling single-cell levels (< 200, 000 counts; (Ding et al., 2020; Jiang et al., 2022; Ziegenhain et al., 2017)). For example, bayesReact tends to better differentiate between samples with and without miR-122-5p expression even for extremely sparse count data (Fig. 4C, Supplementary Fig. S6). Particularly, the medians of the miR-122-5p activity in non-LIHC cancer types are closer to zero for bayesReact compared to miReact (Supplementary Fig. S6). These results indicate an improved performance over miReact on sequencing data with high zero-count content and may subsequently indicate an improved performance on scRNA-Seq data.

### 3.2. Recovering miRNA activities at the single-cell level

To evaluate the performance of bayesReact further, we inferred the miRNA activity in a whole-transcriptome, low-throughput scRNA-Seq dataset (Isakova et al., 2021). We considered the expression and activity of 225 miRNAs across four time points for mESCs differentiating from pluripotent stem cells (day 0) into embryoid bodies (day 12), which contain three germ layers of multipotent and lineage-committed stem cells. All miRNAs, except miR-298-5p, have a mean expression < 2 TPM, and only ten cases are expressed in more than 10% of the cells (> 91 of 913 cells; Supplementary Fig. S7A). The high zero count in the expression data entails both limited miRNA detection and resolution of the 3’ UTR sequence ranks, potentially explaining the reduced correlation scores compared to the bulk TCGA data (Fig. 3A, Fig. 6A).

**Figure 5.**
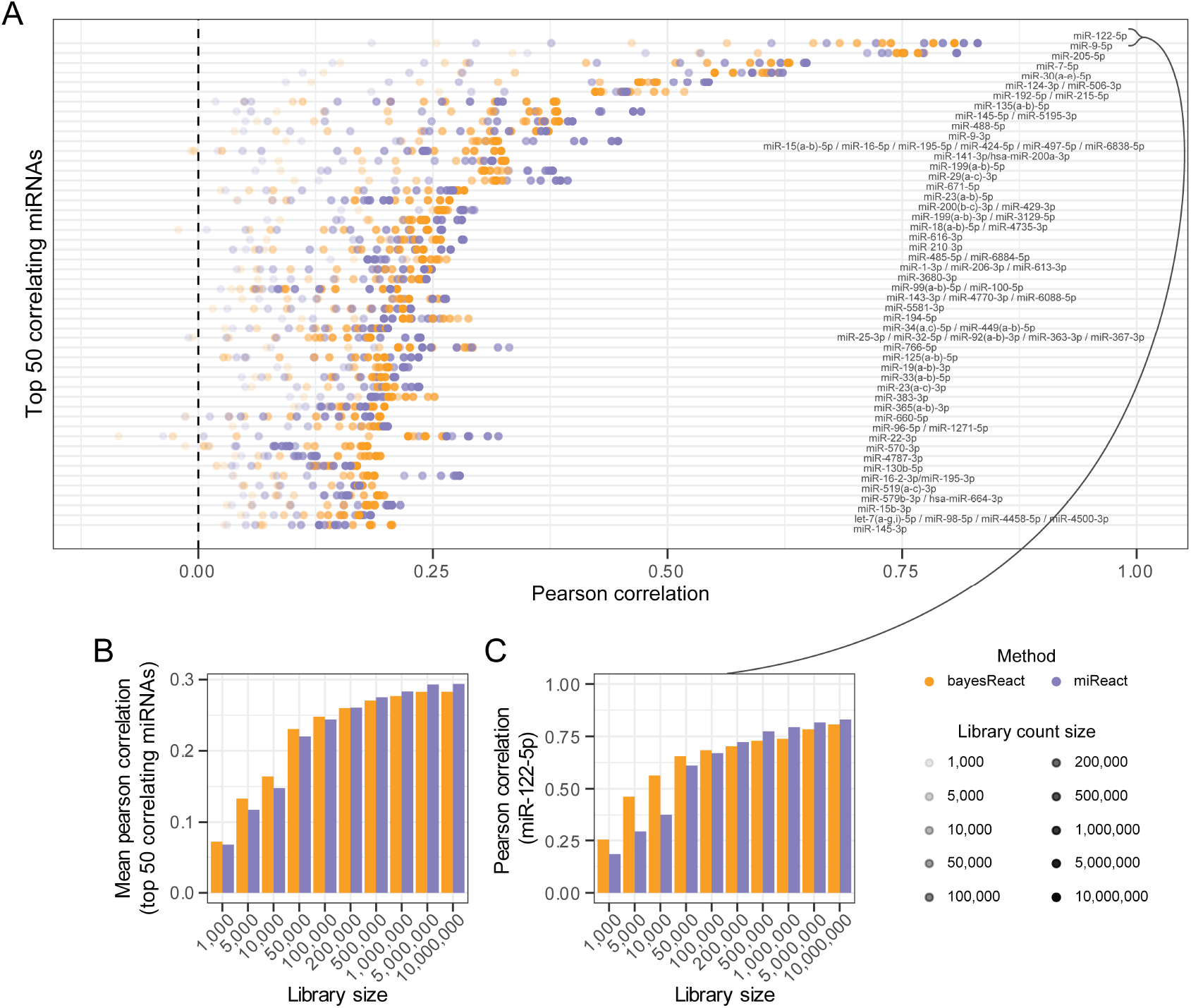
Pan-cancer miRNA activity and expression correlation for varying library sizes, created through downsampling. (A) Pancancer Pearson correlation between miRNA activity and expression for library sizes of different count sparsity. Activities were inferred through bayesReact (orange) and miReact (purple). miRNAs match Fig. 3 panel C. (B) Mean Pearson correlation for the top 50 correlating miRNAs at differing count sparsity. (C) miR-122-5p correlation for varying library size.

**Figure 6.**
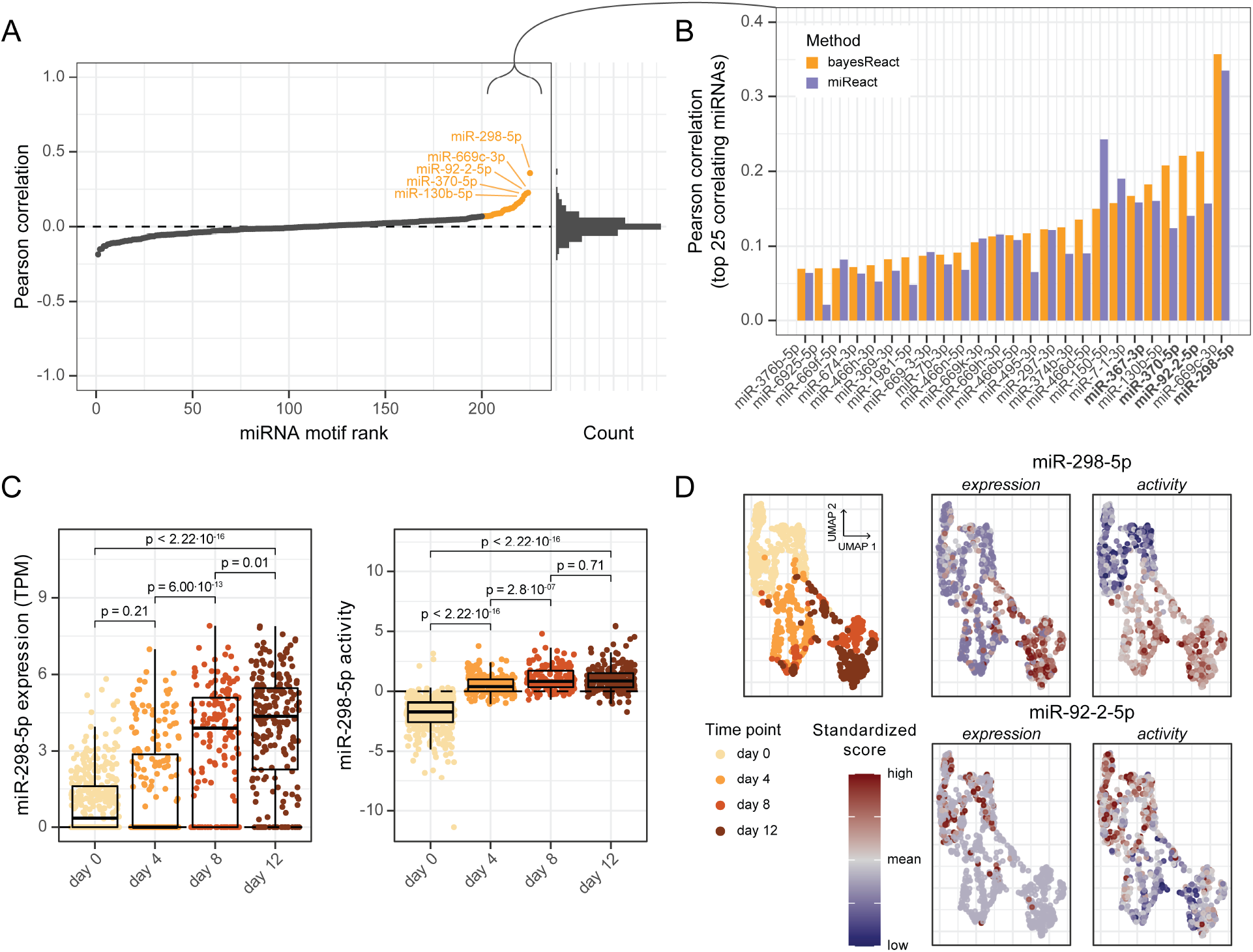
Recovering miRNA activities at the single-cell level. (A) Pearson correlation between miRNA activity and expression across all cells (n = 913), ranked by their correlation coefficient. miRNA activities are obtained using bayesReact. The corresponding histogram, containing 50 bins, is shown on the right. The top 25 correlating miRNAs are highlighted with orange, and the top 5 are annotated. (B) Comparison of correlation coefficients of top 25 miRNAs for activities inferred through bayesReact and miReact. miRNAs highlighted by Isakova et al. are shown in bold. (C) Boxplot with underlying data points depicting miR-298-5p expression (left) and activity (right) across four time points, measured in days. P-values are obtained from Wilcoxon rank-sum tests. TPM = transcripts per million. (D) Cell clustering based on UMAP coordinates. Cells are annotated by time point (left), miRNA expression (middle), and miRNA activity (right). The expression and activity are standardized for visualization purposes.

Focusing on the top 25 correlating miRNAs (Fig. 6B), bayesReact recovers the activity of several miRNAs reported by Isakova et al. as having a temporal expression pattern. This includes miR-298-5p found to have increasing expression and activity during cell differentiation (Fig. 6C-D, Supplementary Fig. S7B), and miR-92-2-5p having decreased activity over time (Fig. 6D, Supplementary Fig. S7D-F). Additionally, miR-370-5p and miR-367-3p showed agreement between expression and activity (Supplementary Fig. S7H). In all instances, we observe a miRNA expression dropout for a subset of cells while the expression is moderate to high for the remaining cells at the same time point (Fig. 6C, Supplementary Fig. S7D,H). In contrast, the activity score does not exhibit a similar zero inflation (Fig. 6C, Supplementary Fig. S7E,H). A zero count can either be biological (no transcript present) or non-biological (failure to measure the transcript) (Jiang et al., 2022; Kharchenko et al., 2014). Since the inferred activity depends on a motif signal relying on the expression-based ranking of all protein-coding transcripts, we suggest that it may be more robust to non-biological dropout. This is consistent with bayesReact continuing to recover the miR-122-5p liver-specific activity from increasingly small library sizes (subsection 3.1.2).

Comparing bayesReact to miReact, we find that bayesReact produces a higher correlation between miRNA activity and expression for 76% of the top 25 miRNAs, with the methods having a mean Pearson correlation of 0.13 and 0.11, respectively (Fig. 6B). Both methods recover an increase in the miR-298-5p activity over time, which correlates with the observed expression (*ρ*_*p*_ = 0.36 for bayesReact, and *ρ*_*p*_ = 0.34 using miReact; Fig. 6C, Supplementary Fig. S7C). However, only bayesReact efficiently recovers the decrease in miR-92-2-5p activity during embryonic stem cell differentiation (*ρ*_*p*_ = 0.22 for bayesReact, and *ρ*_*p*_ = 0.14 with miReact; Supplementary Fig. S7D-G). Furthermore, bayesReact most effectively captures the same significant differences observed for the miRNA expression between time points (Fig. 6C, Supplementary Fig. S7C,D-E,G).

Investigating the top correlating miRNAs further, we find that the miR-297-669 cluster (including miR-297, miR-466, and miR-669) comprises many of the top correlating miRNAs (10 of 25, where only miR-669h-3p and miR-669k-3p share a seed site). The miRNA cluster is derived from an intronic locus in the Sfmbt2 gene (Lehnert et al., 2011), which has previously been shown to be expressed in mouse embryonic stem cells (Zheng et al., 2011; Inoue et al., 2017). Similarly, we observe miR-669c-3p expression in several of the mESCs from day 0 and positive activity for most cells at this time point (Supplementary Fig. S7H). The miRNAs from this cluster have been implicated in developmental, apoptotic, and toxic response processes (Zheng et al., 2011; Inoue et al., 2017; Druz et al., 2012; Liu and Bain, 2018), and our findings further support the potential involvement of miRNAs from the miR-297-669 cluster in murine developmental processes.

## 4. Discussion

We present a new tool for regulatory motif activity inference, which can recover miRNA activities at the bulk and single-cell levels.

bayesReact currently uses a pseudo-normal condition for FC-based 3’ UTR ranking, defined as the median expression of genes across all conditions, e.g., the TCGA samples. Subsequently, the 3’ UTR rank and location along the combined sequence interval are expected to be driven by transcriptional differences between tissues and cancer types. We find that bayesReact can efficiently recover the activity of miRNAs in cancer types where the corresponding miRNA is also expressed. However, when investigating regulatory perturbations during tumorigenesis, we must compare healthy and disease conditions (Rasmussen et al., 2013). This still remains challenging due to the limited access to healthy control samples.

We report a strong negative association between the miR-124-3p activity and REST expression in low-grade gliomas, agreeing with the known negative feedback loop in which both miR-124-3p and the REST protein complex participate to regulate the neuronal cell fate (Visvanathan et al., 2007; Suster and Feng, 2021). Both miR-124-3p down-regulation and REST up-regulation have been implicated in the development and prognosis of gliomas, including the gain of stem-like features, e.g., self-renewal, and tumor invasiveness (Silber et al., 2008; Jiang et al., 2016; Bhaskaran et al., 2019; Conti et al., 2012; Wang et al., 2023a; Xia et al., 2012; Ferrarese et al., 2014; Kamal et al., 2012). However, limited knowledge exists regarding the joint deregulation of the regulatory miR-124-3p/REST axis in cancer (Conti et al., 2012). We propose that miR-124-3p down-regulation in gliomas alleviates the repression of REST, which would further implicate its tumor-suppressor capabilities and therapeutic potential. In the future, inference of healthy control samples may further help us evaluate the perturbation of the miR-124-3p activity and REST expression in the TCGA low-grade glioma samples.

Our results highlight that miRNA activity inference may contribute information even when the miRNA expression is observed, e.g., by elucidating differences in the degree a miRNA is acting on its targets or detecting potential dropout events in whole-transcriptome scRNA-Seq data. For example, Isakova et al. did not originally report on the miR-297-669 cluster, which may be due to its low expression in their data. However, the combined expression and activity of miR-669c-3p, from the miRNA cluster, indicate its presence in the earliest stage of murine stem-cell differentiation, concordant with the known role of the miR-297-669 cluster in developmental processes (Zheng et al., 2011; Inoue et al., 2017). A key prospect of mRNA-based regulatory inference is the use cases for poly(dA)-selected RNA-Seq data, which includes most scRNA-Seq data to date (Wang et al., 2023b). bayesReact is an unsupervised method, allowing for direct miRNA activity inference from high-throughput scRNA-Seq experiments, where miRNA expression is unavailable. The model can potentially be further optimized for sparse count data by letting *µ* (Fig. 2E) be cluster-specific, e.g., allowing the model to learn tissue or cell-type patterns for the regulatory motif distributions.

During this study, we only explored the activity inference of miRNAs, as they are highly studied and well-characterized, making them ideal for model validation. However, many other regulators promote cell homeostasis through motif interaction (Van Roey and Davey, 2015; Lambert et al., 2018; Corley et al., 2020; Liu and Chen, 2022). In a recent study, we used miReact for de-novo motif discovery, which led to the finding of a circHIPK3/IGF2BP2 regulatory axis (Okholm et al., 2024), depicting the use of activity inference for RBPs. Similarly, bayesReact presents a generic hypothesis-generating tool that can also be used for large data screens. It may help detect active motif-based regulatory mechanisms and perform de-novo motif discovery, subsequently identifying candidates for further computational and experimental validation. In addition, we focus on expression-coupled motif modeling using RNA-Seq data for sequence ranking, but other continuous experimental settings could also be used. Advancements in high-throughput proteomics (Cui et al., 2022; Messner et al., 2023) and image-based spatial transcriptomics (Wang et al., 2023c) represent anticipated novel avenues.

In conclusion, we introduce the versatile hypothesis-generating tool bayesReact, which improves the ability to infer condition-specific regulatory motif activity at the single-cell level. The unsupervised method enables comprehensive data screens for regulatory activity detection and provides uncertainty evaluation for activity estimates. We show that it recovers miRNA activities with positive associations to the corresponding measured expressions in bulk and single-cell RNA-Seq data. Furthermore, bayesReact outperforms its predecessor, miReact, on sparse count data, recovering significant temporal miRNA activity patterns in agreement with observed miRNA expression and current literature. We highlight the ability of activity inference to contribute additional information to the miRNA expression by shedding light on the association between the miRNA activity and known target expression and potentially being more robust to dropout events in Smart-seq-total data. In the future, the proposed generative process at the core of bayesReact can be extended, e.g., to explore the changes in target efficiency across cells and other conditions.

## 5. Acknowledgements

Most of the computing was performed on the GenomeDK cluster, and we thank the GenomeDK team and Aarhus University for providing computational resources and support.

We also thank Morten Muhlig Nielsen for helpful discussions relating to data processing and normalization.

## 6. Conflict of interest

None declared.

## 7. Author contributions statement

J.S.P. and A.B.C. conceived and supervised the project and the manuscript preparation. A.B.C. and J.S.P. conceived the probabilistic model with additional contributions from A.M.R. A.M.R. acquired and processed the data, implemented bayesReact, evaluated the model, and analyzed the data. A.M.R. generated the figures and drafted the manuscript.

## 8. Funding

This study was supported with funding from the Novo Nordic Foundation [NNF18OC0053222], Harboe Foundation [2023-0285], and NEYE Foundation (received 07/09/2023).

## 9. Data availability

All data used during this study is publicly available. Matched mRNA and miRNA expression data from primary tumors were obtained from the Cancer Genome Atlas through the Recount3 project and GDC data portal (Weinstein et al., 2013; Wilks et al., 2021; Grossman et al., 2016). Whole-transcriptome, single-cell Smart-seq-total data was obtained from Isakova et al. (Isakova et al., 2021), through the Gene Expression Omnibus (GEO) using the accession number GSE151334.

## Notes

### Competing Interest Statement

The authors have declared no competing interest.

