## Supplementary material for "bayesReact: Expression-coupled regulatory motif analysis detects microRNA activity in cancer and at the single cell level"

doi: NA

### bayesReact: Supplementary material

[1]Department of Clinical Medicine, Aarhus University, Palle Juul-Jensens Boulevard 11, 8200, Aarhus N, Denmark, [2]Department of Molecular Medicine, Aarhus University Hospital, Palle Juul-Jensens Boulevard 11, 8200, Aarhus N, Denmark, [3]Department of Statistics, University of British Columbia, 2207 Main Mall, V6T 1Z4, British Columbia, Canada and [4]Bioinformatics Research Center, Aarhus University, Universitetsbyen 81, 8000, Aarhus C, Denmark

[†]The authors wish it to be known that, in their opinion, the last two authors should be regarded as Joint Last Authors.

#### 1. Supplementary Figures

This section contains the following supplementary figures:

- **Supplementary Figure 1.** Overview of bayesReact input processing.
- **Supplementary Figure 2.** bayesReact content.
- **Supplementary Figure 3.** Model diagnostics and evaluation.
- **Supplementary Figure 4.** Heatmaps depicting clustering of pan-cancer samples.
- **Supplementary Figure 5.** Tissue-specific miR-122-5p and miR-124-3p activities.
- **Supplementary Figure 6.** miR-122-5p activity inference based on differing degrees of library count down-sampling.
- **Supplementary Figure 7.** Recovering miRNA activities at the single-cell level.

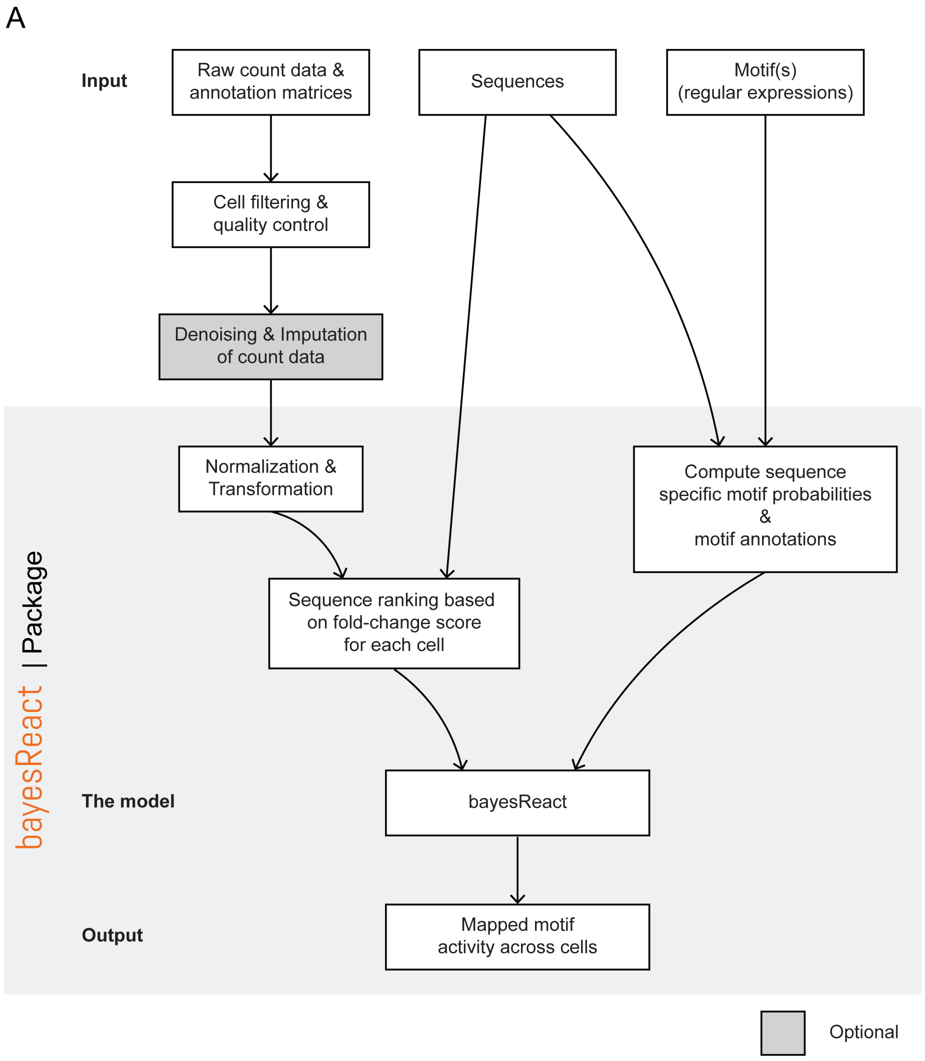


**Supplementary Figure 1.** Overview of bayesReact input processing. (A) bayesReact takes as input filtered expression data, a set of sequences, and regulatory motifs of interest. After manual preprocessing, the bayesReact package handles expression data normalization, sequence ranking, motif annotations, and modeling (highlighted with a grey background). Edges show the flow of data between each processing step.

A

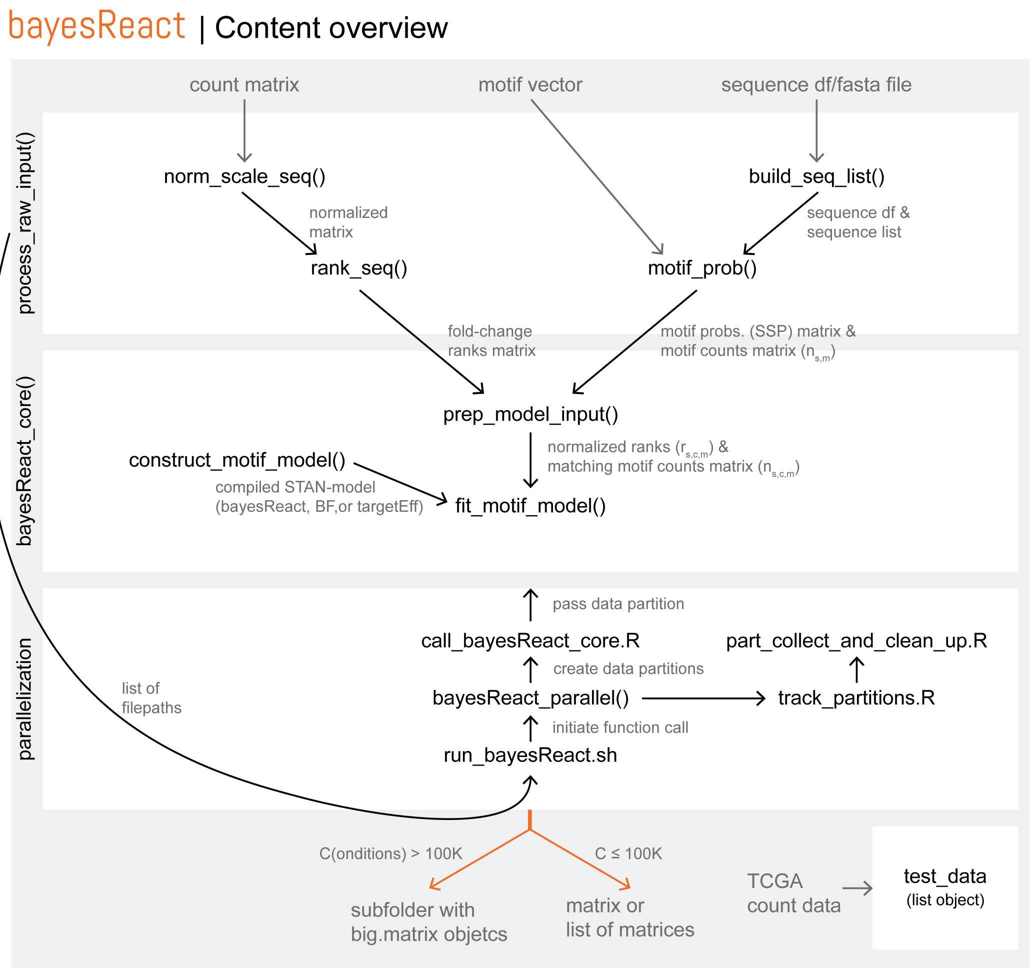


**Supplementary Figure 2.** bayesReact content. (A) Overview of functions and scripts provided in the R-package (black) and the data input (grey), with edges depicting the data flow between function calls. bayesReact contains three main components (white boxes): process_raw_input(), bayesReact_core(), and bayesReact_parallel(). 100K = 100,000.

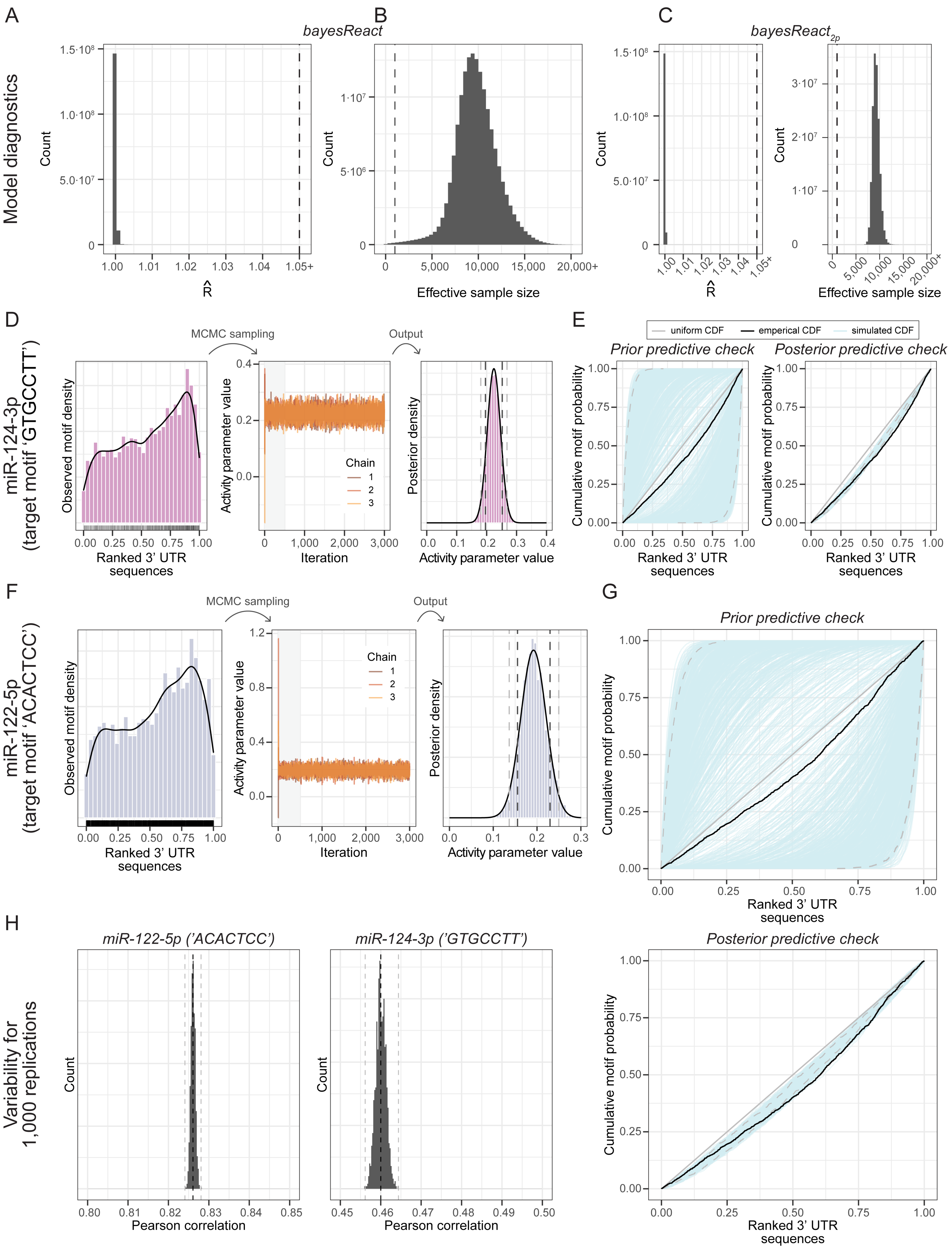

A
B
C
bayesReact
bayesReact2p
Model diagnostics
Count
R̂
Effective sample size
1.00
1.01
1.02
1.03
1.04
1.05+
0
5,000
10,000
15,000
20,000+
D
miR-124-3p
(target motif 'GTGCCTT')
MCMC sampling
Output
Observed motif density
Activity parameter value
Posterior density
Chain
1
2
3
Ranked 3' UTR sequences
Iteration
E
uniform CDF
emperical CDF
simulated CDF
Prior predictive check
Posterior predictive check
Cumulative motif probability
F
miR-122-5p
(target motif 'ACACTCC')
G
H
Variability for
1,000 replications
miR-122-5p ('ACACTCC')
miR-124-3p ('GTGCCTT')
Pearson correlation

**Supplementary Figure 3.** Model diagnostics and evaluation. (A) MCMC chain convergence diagnostic ($\hat{R}$) for all bayesReact activity parameters for the TCGA data ($n = M \times C \approx 158 \cdot 10^6$). $\hat{R}$ compares the within and between chain parameter estimates, with values close to one supporting convergence on the stationary distribution (posterior). The dashed line highlights $\hat{R} = 1.05$, and the histogram contains 50 bins, with values capped at 1.05+. (B) Effective sample size (ESS) for all activity parameters from the TCGA data. ESS is an MCMC chain auto-correlation diagnostic and a small ESS indicates high uncertainty of a parameter estimate due to inefficient exploration of the stationary target distribution. The dashed line highlights $ESS = 1,000$ and the histogram contains 50 bins, with values capped at 20,000+. (C) Diagnostics plots for the two-parameter beta model (bayesReact$_{2p}$), equivalent to panel A-B. The plots depict $\hat{R}$ and $ESS$ for each $\tau_{c,m}$ parameter. (D) Overview of the marginal posterior approximation process for a single activity parameter, $a_{7278,11872}$, parameterizing the miR-124-3p target motif distribution across sequences ranked by fold-change scores from a low-grade glioma (LGG) sample. Left: The observed motif distribution across the normalized combined sequence interval of all 3' UTRs. The histogram contains 30 bins, and a corresponding density line is shown, while the rug (bottom) depicts the underlying motif observations represented by the midpoint, $r_{i,7278,11872}$, of a given sequence. Middle: Traceplot of three MCMC chains run for 3,000 iterations, including a discarded warm-up period of 500 iterations (grey shading). Right: The MCMC approximation of the marginal posterior (histogram with 50 bins). The normal approximation (black line) is defined by the posterior mean and standard deviation and is used to find the motif activity. The dashed lines show the 80% (dark) and 95% (light) credible intervals (CI). (E) Cumulative motif distributions sampled from the prior (right) and posterior (left) predictive distributions. The observed empirical cumulative distribution function (CDF; black) and theoretical uniform CDF (grey) are plotted together with 1,000 simulated CDFs (blue). A thousand $a_{7278,11872}$ values were randomly sampled from the prior and posterior distributions. Predictive CDFs were then generated by simulating motif occurrences under beta distributions parameterized by each sampled activity parameter, with sample sizes equal to the total number of motif observations ($n_m = 3,067$). The dashed lines depict the predictive CDFs of the 95% CI for the sampled activity parameters. (F) The marginal posterior approximation process for $a_{406,1142}$, which parameterizes the miR-122-5p target motif distribution in a liver hepatocellular carcinoma (LIHC) sample. Plots corresponding to panel D are depicted, showing the observed motif distribution (left), traceplot (middle), and marginal posterior distribution (right). (G) Prior (top) and posterior (bottom) predictive checks for $a_{406,1142}$, equivalent to analysis shown in panel E, with $n_m = 1,692$. (H) Pan-cancer Pearson correlation between miRNA expression and activity for a 1,000 replications of the bayesReact activity inference for the miR-122-5p activity (left) and miR-124-3p activity (right). The dashed lines highlight the median correlation (dark) as well as the minimum and maximum values (light). The histograms contain 50 bins.

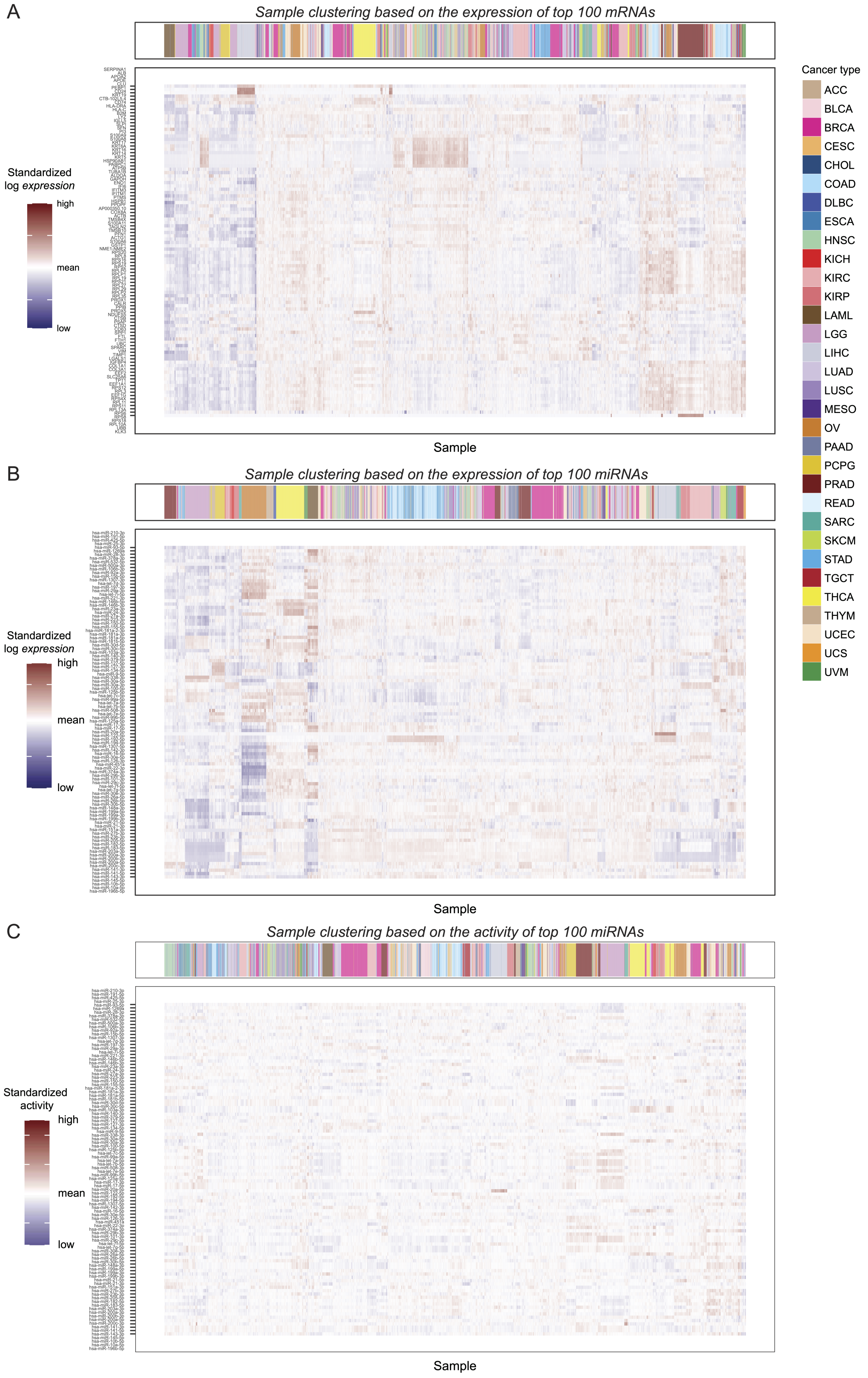
A
Sample clustering based on the expression of top 100 mRNAs
Standardized log expression
high
mean
low
Sample
B
Sample clustering based on the expression of top 100 miRNAs
Standardized log expression
high
mean
low
Sample
C
Sample clustering based on the activity of top 100 miRNAs
Standardized activity
high
mean
low
Sample
Cancer type
ACC
BLCA
BRCA
CESC
CHOL
COAD
DLBC
ESCA
HNSC
KICH
KIRC
KIRP
LAML
LGG
LIHC
LUAD
LUSC
MESO
OV
PAAD
PCPG
PRAD
READ
SARC
SKCM
STAD
TGCT
THCA
THYM
UCEC
UCS
UVM

**Supplementary Figure 4.** Heatmaps depicting clustering of pan-cancer samples. (A) Heatmap of the log-transformed mRNA expression across all TCGA samples for the 100 transcripts with the highest mean RPKM values. Hierarchical agglomerative clustering (HAC) was performed on both the transcript and sample level, with dissimilarities between observations measured using Euclidean distances and complete linkage used to evaluate distances between sets of observations. Standardized expression values are used for visualization. RPKM = reads per kilobase of transcript per million mapped reads. (B) Heatmap depicting top 100 miRNAs based on highest mean log-transformed TPM values. HAC was performed on both the miRNA transcript and sample level, and standardized values are used for visualization. TPM = transcripts per million. (C) Heatmap showing the standardized activity of miRNAs from panel B. Independent HAC was performed at the sample level.

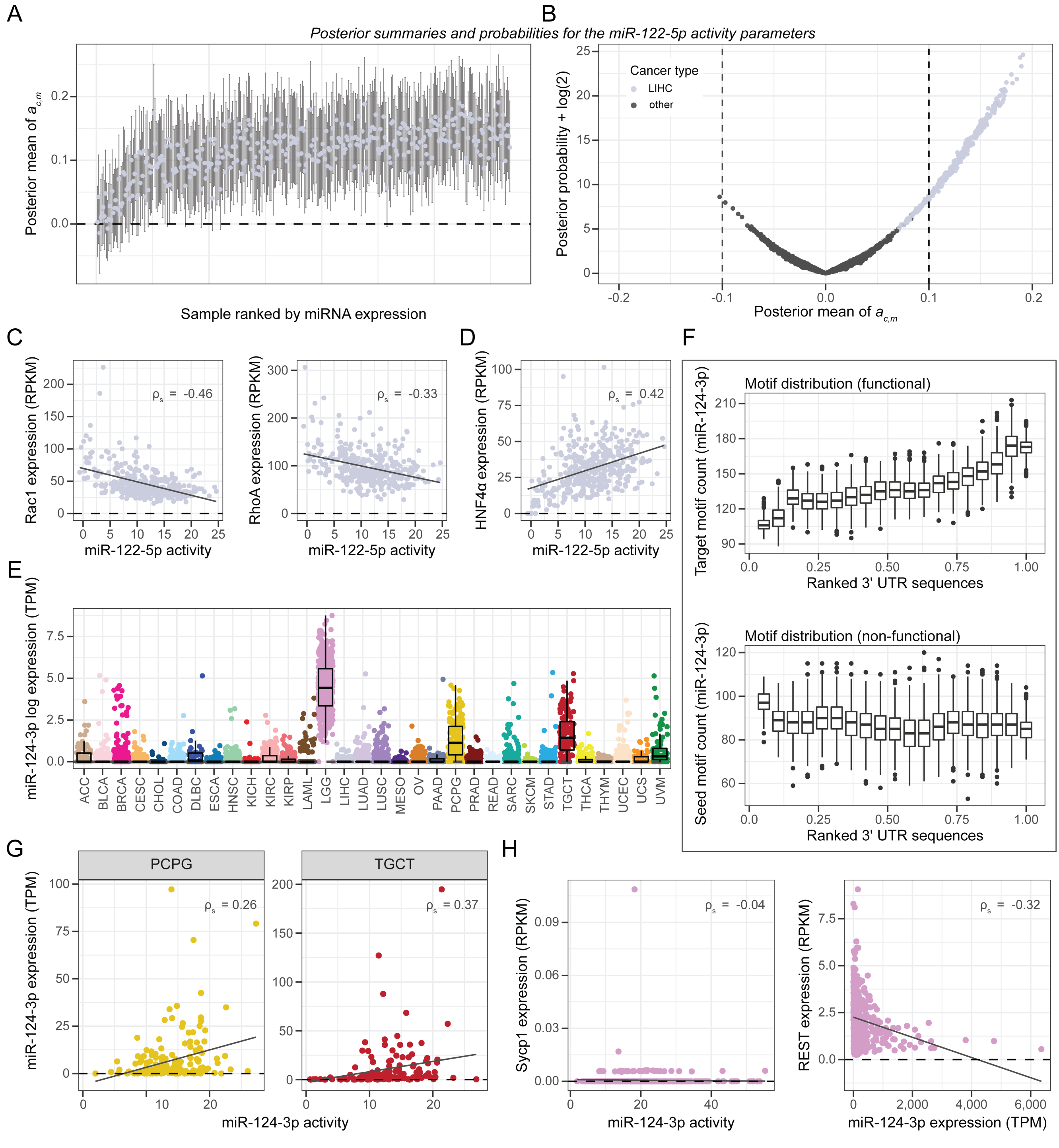


**Supplementary Figure 5.** Tissue-specific miR-122-5p and miR-124-3p activities. (A) The (marginal) posterior means and 99% credible intervals (CIs) for the activity parameter $a_{c,m}$ of the miR-122-5p target motif across all liver hepatocellular carcinoma (LIHC) samples (n = 367). Samples are ordered by observed miR-122-5p normalized expression. (B) Posterior mean for the miR-122-5p activity parameter from each TCGA sample plotted against the corresponding posterior probability of $\mathbf{1}[\bar{a}_{c,m} \geq 0] \log P(a_{c,m} \leq 0 \mid \mathbf{n_{c,m}}) + \mathbf{1}[\bar{a}_{c,m} < 0] \log P(a_{c,m} \geq 0 \mid \mathbf{n_{c,m}}) + \log(2)$, where $\bar{a}_{c,m}$ is the posterior mean. (C) The miR-122-5p activity plotted against the expression of two target genes, Rac1 (left) and RhoA (right), for the LIHC samples. (D) The miR-122-5p activity plotted against the expression of HNF4$\alpha$, which is a transcription factor promoting the transcription of the miR-122 host gene. (E) Log-transformed miR-124-3p expression across all primary tumor samples from the TCGA data. One is added to the expression provided as transcripts per million (TPM + 1). (F) miR-124-3p target site (top) and seed site (bottom) distribution across the normalized 3' UTR sequences. The combined sequence interval is divided into 20 bins, and each boxplot depicts the motif count within a bin for each low-grade glioma (LGG) sample (n = 509). The miR-124-3p target motif occurs 3,067 times across 2,624 3' UTRs, while the seed motif occurs 1,824 times in 1,661 3' UTRs. (G) The miR-124-3p activity plotted against its expression in the pheochromocytomas and paragangliomas (PCPG) samples (n = 182; left) and testicular germ-cell tumors (TGCT) samples (n = 139; right). A linear regression line and Spearman correlation are shown. (H) The miR-124-3p activity plotted against the expression of its known target Sycp1 (left), and the miR-124-3p expression plotted against its downstream target REST (right) for the low-grade glioma samples. A linear regression line and Spearman correlation are depicted. RPKM = reads per kilobase of transcript per million mapped reads; TPM = transcripts per million; UTR = untranslated region.

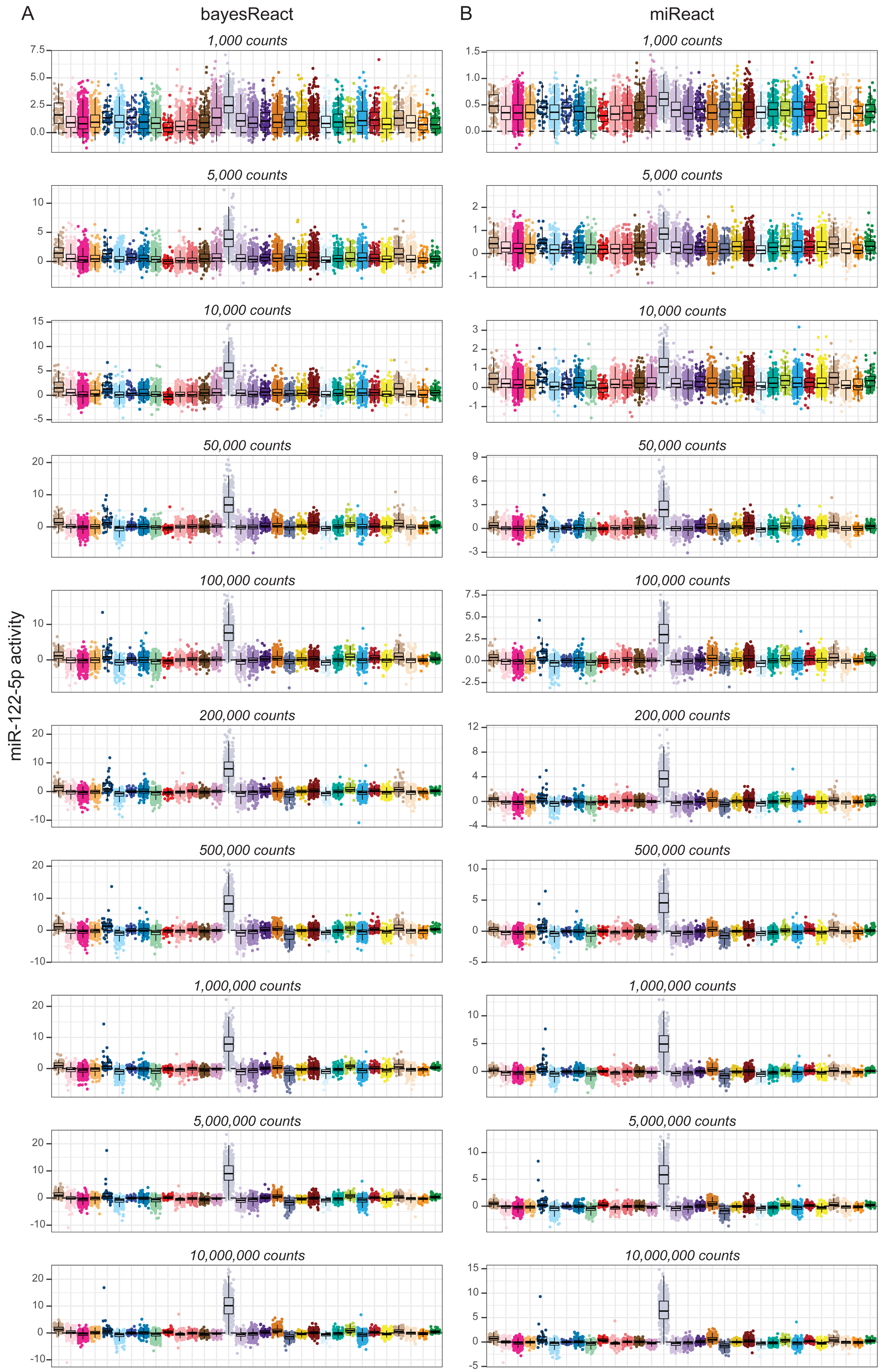
A
bayesReact
B
miReact
1,000 counts
5,000 counts
10,000 counts
50,000 counts
100,000 counts
200,000 counts
500,000 counts
1,000,000 counts
5,000,000 counts
10,000,000 counts
miR-122-5p activity

**Supplementary Figure 6.** miR-122-5p activity inference based on differing degrees of library count down-sampling. (A) miR-122-5p activity inference using bayesReact with each row showing results for increasing library count sizes. The cancer type order is the same as Figure 4 panel A, with liver hepatocellular carcinoma (LIHC) samples shown in grey. (B) Corresponding miR-122-5p activity inference using miReact.

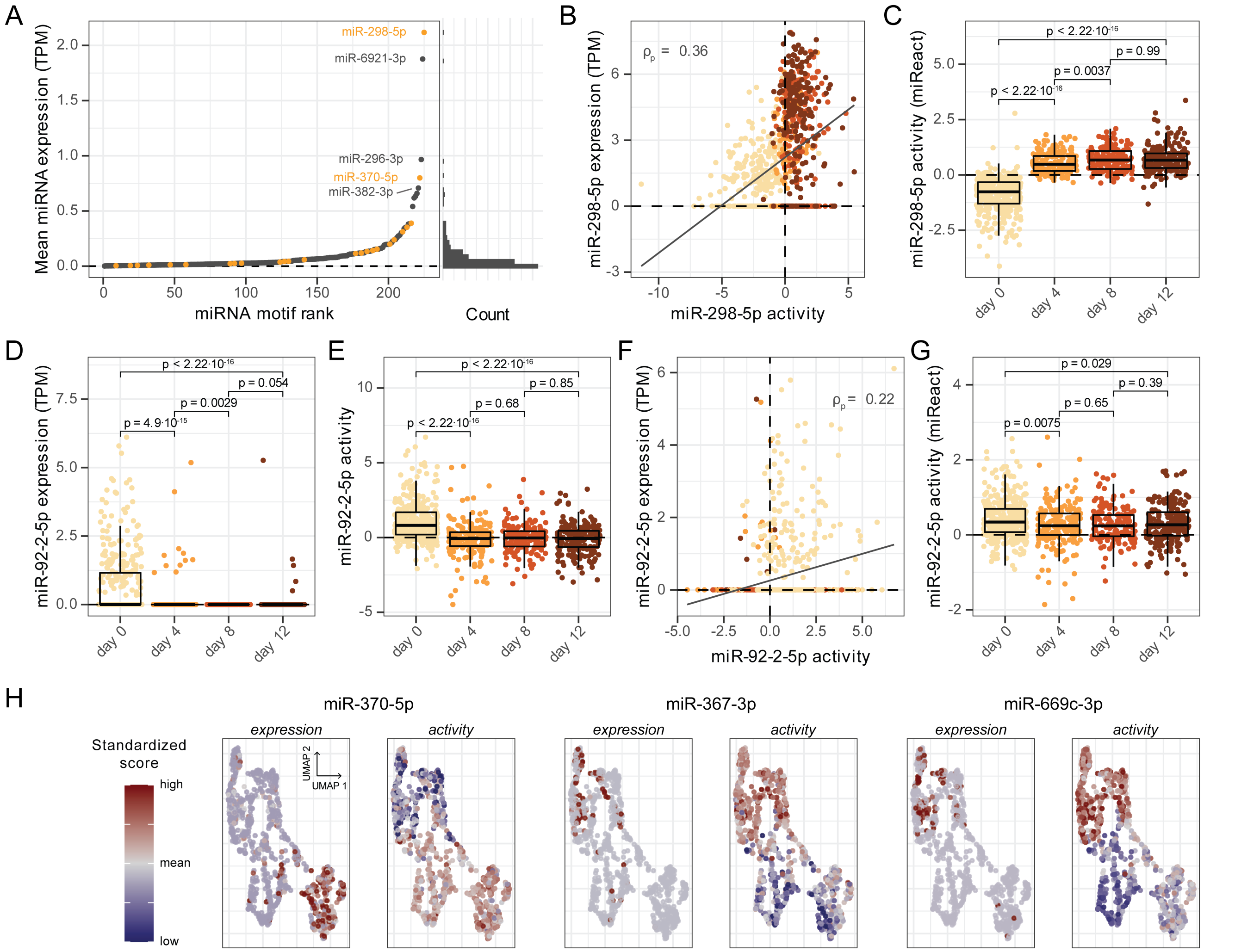


**Supplementary Figure 7.** Recovering miRNA activities at the single-cell level. (A) Mean miRNA expression across all cells ordered by the mean values (left) and corresponding histogram containing 50 bins (right). Top 25 miRNAs with the highest correlation between expression and activity are highlighted in orange. TPM = transcripts per million. (B) miR-298-5p activity plotted against the expression and linear regression line depicted. Points are colored by sample extraction time. TPM = transcripts per million. (C) miR-298-5p activity obtained through miReact depicted per time point. Wilcoxon rank-sum test was performed and annotated for each compared time point. (D) miR-92-2-5p expression over time. P-values were obtained from Wilcoxon rank-sum tests. TPM = transcripts per million. (E) Boxplots depicting the miR-92-2-5p activity over time. P-values were obtained from Wilcoxon rank-sum tests. (F) The miR-92-2-5p activity plotted against the observed expression, and linear regression line shown. Points are colored by sample extraction time. (G) The miR-92-2-5p activity obtained through miReact is depicted per time point. P-values were obtained from Wilcoxon rank-sum tests. (H) Dimension reduction plots based on UMAP coordinates depicting cell clustering, with expression and activity annotated. The expression and activity are standardized to have a mean of zero and a standard deviation of one. Please see Figure 6 panel D for time point annotations.

#### 2. Supplementary Tables

This section contains the following supplementary tables:

- **Supplementary Table 1.** Overview of cancer types in the TCGA data.
- **Supplementary Table 2.** Names and target sites of top 50 correlating miRNAs from the TCGA data.

| Cancer type | Cancer name | Primary site | # samples |
|---|---|---|---|
| BRCA | Breast Invasive Carcinoma | Breast | 1068 |
| UCEC | Uterine Corpus Endometrial Carcinoma | Uterus | 530 |
| LGG | Brain Lower Grade Glioma | Brain | 509 |
| THCA | Thyroid Carcinoma | Thyroid | 504 |
| LUAD | Lung Adenocarcinoma | Lung | 503 |
| KIRC | Kidney Renal Clear Cell Carcinoma | Kidney | 502 |
| HNSC | Head and Neck Squamous Cell Carcinoma | Head and Neck | 497 |
| PRAD | Prostate Adenocarcinoma | Prostate | 493 |
| LUSC | Lung Squamous Cell Carcinoma | Lung | 475 |
| COAD | Colon Adenocarcinoma | Colorectal | 434 |
| OV | Ovarian Serous Cystadenocarcinoma | Ovary | 419 |
| STAD | Stomach Adenocarcinoma | Stomach | 408 |
| BLCA | Bladder Urothelial Carcinoma | Bladder | 403 |
| LIHC | Liver Hepatocellular Carcinoma | Liver | 367 |
| CESC | Cervical Squamous Cell Carcinoma and Endocervical Adenocarcinoma | Cervix | 304 |
| KIRP | Kidney Renal Papillary Cell Carcinoma | Kidney | 291 |
| SARC | Sarcoma | Soft Tissue | 257 |
| ESCA | Esophageal Carcinoma | Esophagus | 183 |
| PCPG | Pheochromocytoma and Paraganglioma | Adrenal Gland | 182 |
| PAAD | Pancreatic Adenocarcinoma | Pancreas | 178 |
| LAML | Acute Myeloid Leukemia | Bone Marrow | 173 |
| READ | Rectum Adenocarcinoma | Colorectal | 159 |
| TGCT | Testicular Germ Cell Tumors | Testis | 139 |
| THYM | Thymoma | Thymus | 120 |
| SKCM | Skin Cutaneous Melanoma | Skin | 97 |
| MESO | Mesothelioma | Pleura | 87 |
| UVM | Uveal Melanoma | Eye | 80 |
| ACC | Adrenocortical Carcinoma | Adrenal Gland | 79 |
| KICH | Kidney Chromophobe | Kidney | 66 |
| UCS | Uterine Carcinosarcoma | Uterus | 57 |
| DLBC | Lymphoid Neoplasm Diffuse Large B−cell Lymphoma | Lymph Nodes | 47 |
| CHOL | Cholangiocarcinoma | Bile Duct | 36 |

**Supplementary Table 1.** Overview of cancer types in the TCGA data. The table depicts all cancer types included from the TCGA data, and contains the abbreviated cancer type (column 1); full name of the cancer type (column 2); tissue origin of the primary tumor (column 3); and number of samples assigned to each cancer type (column 4).

| miRNA | # miRNAs sharing target site | Target site |
|---|---|---|
| miR−122−5p | 1 | ACACTCC |
| miR−9−5p | 1 | ACCAAAG |
| miR−205−5p | 1 | ATGAAGG |
| miR−7−5p | 1 | GTCTTCC |
| miR−30a−5p/miR−30b−5p/miR−30c−5p/miR−30d−5p/miR−30e−5p | 5 | TGTTTAC |
| miR−124−3p/miR−506−3p | 2 | GTGCCTT |
| miR−192−5p/miR−215−5p | 2 | TAGGTCA |
| miR−135a−5p/miR−135b−5p | 2 | AAGCCAT |
| miR−145−5p/miR−5195−3p | 2 | AACTGGA |
| miR−488−5p | 1 | TATCTGG |
| miR−9−3p | 1 | AGCTTTA |
| miR−15a−5p/miR−15b−5p/miR−16−5p/miR−195−5p/miR−424−5p/miR−497−5p/miR−6838−5p | 7 | TGCTGCT |
| miR−141−3p/miR−200a−3p | 2 | CAGTGTT |
| miR−199a−5p/miR−199b−5p | 2 | ACACTGG |
| miR−29a−3p/miR−29b−3p/miR−29c−3p | 3 | TGGTGCT |
| miR−671−5p | 1 | GGCTTCC |
| miR−23a−5p/miR−23b−5p | 2 | GGAACCC |
| miR−200b−3p/miR−200c−3p/miR−429-3p | 3 | CAGTATT |
| miR−199a−3p/miR−199b−3p/miR−3129−5p | 3 | ACTACTG |
| miR−18a−5p/miR−18b−5p/miR−4735−3p | 3 | GCACCTT |
| miR−616−3p | 1 | CAATGAC |
| miR−210−3p | 1 | ACGCACA |
| miR−485−5p/miR−6884−5p | 2 | CAGCCTC |
| miR−1−3p/miR−206-3p/miR−613-3p | 3 | ACATTCC |
| miR−3680−3p | 1 | ATGCAAA |
| miR−99a−5p/miR−99b−5p/miR−100−5p | 3 | TACGGGT |
| miR−143−3p/miR−4770-3p/miR−6088-5p | 3 | TCATCTC |
| miR−5581−3p | 1 | GCATGGA |
| miR−194−5p | 1 | CTGTTAC |
| miR−34a−5p/miR−34c−5p/miR−449a/miR−449b−5p | 4 | CACTGCC |
| miR−25−3p/miR−32−5p/miR−92a−3p/miR−92b−3p/miR−363−3p/miR−367−3p | 6 | GTGCAAT |
| miR−766−5p | 1 | TTCCTCC |
| miR−125a−5p/miR−125b−5p | 2 | CTCAGGG |
| miR−19a−3p/miR−19b−3p | 2 | TTTGCAC |
| miR−33a−5p/miR−33b−5p | 2 | CAATGCA |
| miR−23a−3p/miR−23b−3p/miR−23c | 3 | AATGTGA |
| miR−383−3p | 1 | AGTGCTG |
| miR−365a−3p/miR−365b−3p | 2 | GGGCATT |
| miR−660−5p | 1 | AATGGGT |
| miR−96−5p/miR−1271−5p | 2 | GTGCCAA |
| miR−22−3p | 1 | GGCAGCT |
| miR−570−3p | 1 | TGTTTTC |
| miR−4787−3p | 1 | GGCGCAT |
| miR−130b−5p | 1 | GAAAGAG |
| miR−16−2−3p/miR−195−3p | 2 | AATATTG |
| miR−519a−3p/miR−519b−3p/miR−519c−3p | 3 | TGCACTT |
| miR−579−3p/miR−664b−3p | 2 | CAAATGA |
| miR−15b−3p | 1 | ATGATTC |
| let−7a−5p/let−7b−5p/let−7c−5p/let−7d−5p/let−7e−5p/let−7f−5p/let−7g−5p/let−7i−5p/miR−98−5p/miR−4458-5p/miR−4500-3p | 11 | CTACCTC |
| miR−145−3p | 1 | AGGAATC |

**Supplementary Table 2.** Names and target sites of top 50 correlating miRNAs from the TCGA data. The table includes the names (column 1) of the top 50 miRNAs with the highest Pearson correlation between expression and activity. The miRNAs are collapsed based on shared target site (columns 2-3).
